# Leveraging the joint site frequency spectrum to detect genomic regions of early divergence

**DOI:** 10.64898/2026.03.03.709333

**Authors:** Alejandro Calderon, Genevieve M. Kozak, Lawrence H. Uricchio

## Abstract

Detecting genomic loci underlying local adaptation is a central goal in population genetics. Such loci have been discovered using a variety of approaches, but the statistical performance of these approaches can depend substantially on the frequencies of the alleles underlying phenotypic adaptation. In particular, multiple evolutionary modeling studies have shown that rare alleles of large effect can make substantial contributions to phenotypic variance and potentially adaptive divergence, but most selection inference methods are not sensitive to rare alleles. We used simulations of evolutionary divergence to compare commonly-used *F_ST_* scans to a likelihood-based approach that interrogates the whole frequency spectrum. We found that the likelihood-based approach outperforms *F_ST_* in simulations in which low frequency alleles play a substantial role in driving trait divergence. We applied both approaches to genomic data from *Ostrinia nubilalis* (the European Corn Borer), a species in which population-specific variation in circannual rhythms and mate preference phenotypes have driven recent reproductive isolation. The likelihood-based approach recovers previously discovered genomic loci and finds several new candidate regions that may be relevant for reproductive isolation between populations. Our findings demonstrate how the two-dimensional frequency spectrum may help to identify loci contributing to reproductive isolation in contexts when commonly used methods are less powerful.

**Article summary:** Identifying genomic loci underlying phenotypic divergence is a fundamental challenge in population genetics. Some commonly used approaches, like *F_ST_* scans, perform poorly when rare alleles drive phenotypic differentiation, a scenario that arises under some polygenic trait architectures. We used forward simulations of selection under different trait architectures to compare *F_ST_* scans to a likelihood-based method, and found that the likelihood-based approach is more powerful when low-frequency alleles contribute to trait divergence. Applying both methods to the European Corn Borer moth, we recovered known loci and identified new candidate divergent regions, demonstrating the utility of likelihood-based approaches in population genomics.

## Introduction

Local adaptation is a major contributor to differentiation between populations, and much of what we know about local adaptation at the genomic level is based on simple genome-wide outlier scans using statistics like *F_ST_* [Beaumont, 2005, Hoban et al., 2016]. These approaches have yielded many candidate adaptive loci across numerous species [Canales-Aguirre et al., 2022, Colli et al., 2014, Defaveri and Merilä, 2013, Funk et al., 2016, Keller et al., 2012], and *F_ST_* remains attractive in many contexts because it is reasonable to apply even when sample sizes are small or when only a draft genome is available – constraints that may make the application of more sophisticated methods infeasible for some non-model organisms. For example, powerful haplotype-based selection scans [Sabeti et al., 2007, Voight et al., 2006] require high quality reference genomes, phased sequences, and sometimes large sample sizes. While recent studies have sought to relax some of these constraints [DeGiorgio and Szpiech, 2022, Harris et al., 2018, Klassmann and Gautier, 2022, Szpiech, 2024], high quality genomes are fundamental to the detection of low-diversity regions under selection, and many non-model species have poorly resolved genomes. Gene-environment associations (GEA) [Lotterhos and Whitlock, 2015] and polygenic selection inference methods [Berg and Coop, 2014, Turchin et al., 2012] require complementary data, such as summary statistics from genome-wide association studies or environmental co-variates, and large samples are frequently required to obtain statistical power. Consequently, *F_ST_* scans have retained utility despite methodological advances.

While *F_ST_* scans have had great success, they have some limitations. Though they are essentially agnostic to the underlying evolutionary process that drives population-specific adaptation, *F_ST_* scans are much more sensitive to alleles that have high frequencies in the pooled sample than alleles with low frequencies, in part because *F_ST_* cannot take large values when alleles are rare [Jakobsson et al., 2013]. Many prior studies have simply ignored low-frequency alleles for this reason [Linck and Battey, 2019]. However, multiple evolutionary modeling studies have shown that large-effect alleles are disproportionately confined to low frequencies under stabilizing selection, because selection against them is far more efficient than against small-effect alleles [Lande, 1975, Uricchio et al., 2019]. Consistent with this, low-frequency alleles have been shown to be contributors to standing phenotypic variation [Simons et al., 2018] and large-effect alleles make disproportionately large per-locus contributions to phenotypic adaptation immediately following a shift in optimal phenotype values, before this contribution declines over longer time scales [Hayward and Sella, 2022]. More generally, the space of possible mechanistic evolutionary models underlying adaptation is vast, and includes selection on standing neutral variation [Prezeworski et al., 2005], beneficial reversal of dominance [Posavi et al., 2014], and selection on recurrent *de novo* mutation [Pennings and Hermisson, 2006], to name a few. Each of these will result in different perturbations to the frequency spectrum, some of which may be easily detectable in the early stages of population divergence using commonly-used selection scans, and some of which may not. It would be desirable to employ statistical inference methods that can detect deviations across the frequency spectrum, especially since low frequency alleles could present hidden reservoirs of adaptive capacity that would be missed by most commonly used approaches.

In principle, if causal alleles for a fitness-related trait are known it is possible to test for evidence of selection acting on these sites to drive phenotypic differences between populations [Berg and Coop, 2014, Turchin et al., 2012]. Unfortunately, rare causal alleles are typically very difficult to detect in standard genome-wide association studies because they must have massive effect sizes to overcome the limitations of statistical power at low frequencies [Hayes, 2013]. Methods have been developed and applied to model species to estimate the genome-wide contribution of rare alleles to trait variance [Hernandez et al., 2019, Wainschtein et al., 2022], in some cases finding substantial variance explained by rare (or even ultra-rare) alleles. If rare alleles do explain substantial trait variance as a class for some traits, then their large per-locus effect sizes and transient allele frequency shifts (explored by Hayward and Sella [2022]) could play a role in adaptation, at least at short time scales after environmental shifts. Currently, we know relatively little about the potential role of large-effect rare alleles in driving trait variance and phenotypic differentiation from an empirical perspective.

If rare alleles do make substantial contributions to trait variance, this pattern is only straightforward to explain through selection acting to constrain alleles with large effects to low frequencies [Eyre-Walker, 2010]. Consequently, rare variants are unlikely to make contributions to variance for traits that are weakly selected or for which effect sizes have only modest variation. Stabilizing selection, which is likely to operate on many environmentally-sensitive traits, will tend to remove alleles with very large effects because individuals carrying such variants tend to be displaced substantially from the optimum trait value [Charlesworth, 2013]. Negative selection and pleiotropy (specifically, pleiotropy in which alleles underlying multiple selected traits have correlated effect sizes) can also drive similar patterns [Johnson and Barton, 2005]. When the environment changes, selection can act to transiently increase the frequencies of very large-effect alleles if their effects align with the direction of the change in the phenotypic optimum. However, these alleles rarely fix in most previously-explored models [Hayward and Sella, 2022], because they rarely obtain high enough frequencies before the population arrives at the new optimum trait value (but see Stetter et al. 2018).

In this study, we hypothesized that a likelihood-based approach might be more sensitive than *F_ST_* scans for some genomic architectures and at some time points in the adaptation process, especially for architectures in which rare alleles explain substantial trait variance at early times in the differentiation process. Likelihood-based approaches have been widely applied to detect directional selection acting across the genome [Boyko et al., 2008, Chen et al., 2010, Keightley and Eyre-Walker, 2007, Racimo and Schraiber, 2014, Sawyer and Hartl, 1992] and to detect unusual distributions of rare alleles at specific genomic loci [Lee et al., 2012, Neale et al., 2011], but have not been widely studied in the context of environmental shifts and polygenic selection. We applied both *F_ST_* scans and a two-population likelihood-based scan (following the approach of sweepFinder, Nielsen et al. 2005, 2009) to simulated genomic data under a range of trait architectures (*i.e.*, traits with differing joint distributions of effect sizes and allele frequencies), and compared the relative statistical performance of each method. We then applied both approaches to genomic data from the European Corn Borer (ECB), a model species for incipient speciation in which some of the loci underlying reproductive isolation have been mapped [Kozak et al., 2019, Kunerth et al., 2022], comparing their performance in an empirical setting. We discuss how likelihood-based approaches could be used to further partition the frequency spectrum to assess the relative contributions of rare alleles to population differentiation, and consider both practical limitations of likelihood-based approaches and methodological opportunities that could improve their utility for the discovery of loci harboring rare variants driving phenotypic differentiation.

## Materials and methods

### Simulations

We used SLiM v4.3 [Haller and Messer, 2023] to generate polymorphism data from two subpopulations undergoing divergent selection (Fig. 1A). To mimic the early stages of a differentiation process, we specifically sought evolutionary scenarios in which high gene flow would occur in the genome background and polygenic selection would result in little difference in the 1D frequency spectrum, but population-specific selection pressures would result in trait divergence (and subtle differences in the 2D spectrum) despite gene flow (Fig. 1).

**Figure 1:**
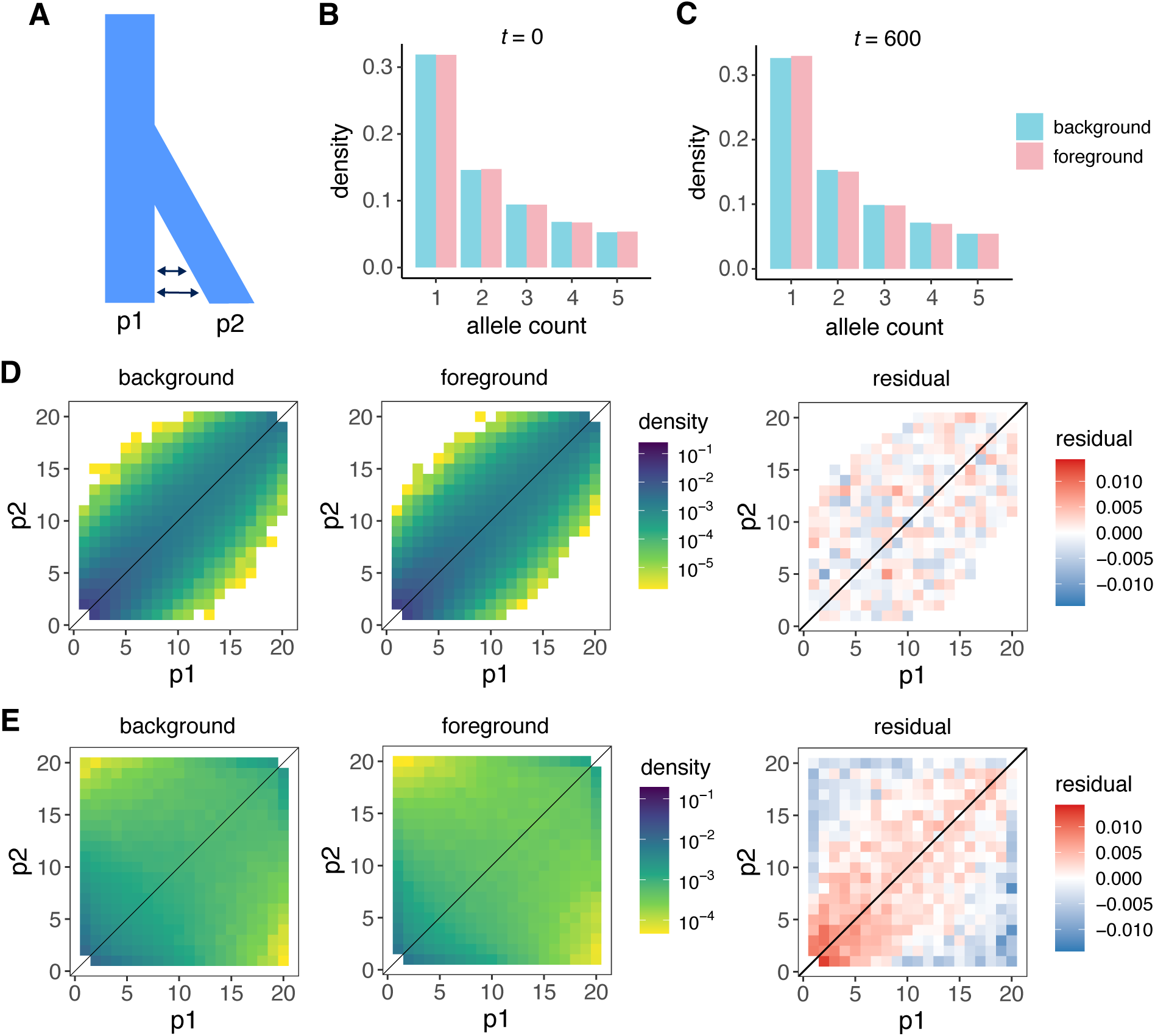
Simulation results pertaining to population split with migration (*m* = 0.01) and divergent selection under the CV model. (**A**). We extracted the 1D frequency spectra for QTLs at the population split (*t* = 0) (**B**) and *t* = 600 (**C**) for foreground loci (undergoing divergent selection) and background loci (undergoing only stabilizing selection) and show the first 5 bins of the spectrum (in a sample of 40 chromosomes). For the same time points, we extracted the 2D spectra for the background and foreground loci, as well as the Poisson residuals between them (background minus foreground, **D** and **E** respectively) (described in Gutenkunst et al. 2009). Red or blue residuals depict an excess or depletion of alleles in a given cell, respectively. There is little difference in the 1D spectrum between the foreground at either time point. In the 2D-spectrum, although the foreground and background frequency spectra are similar, the residuals show a clear deficit of highly differentiated sites in the background at the latter time point.

To accomplish this, we simulated a simple evolutionary model in which an ancestral population splits into two subpopulations. Three distinct quantitative traits are under stabilizing selection. The three traits initially have the same fitness function, but distinct genomic loci drive variation in each trait. We include the distinct quantitative traits for technical reasons – specifically, it would be unfair to assess our ability to detect selection in the divergent locus as compared only to a neutral locus, because the selected locus has a different frequency spectrum even when there is no divergence in the selected trait. We discuss this comparison in more detail in the **Power analyses** section.

After the population split, the optimal trait value gradually increases in one of the subpopulations relative to the ancestral population for one of the three simulated traits, while the fitness function in the other population remains unchanged. For the other traits, there is no change in the fitness function in either population. Though there is high gene flow, alleles may be more likely to increase in frequency in one population if they are beneficial in that context, but this depends on the model parameters and genomic architecture (see next section). Throughout the manuscript, we compare results from two distinct models, one in which common variants explain most heritability (CV model) and another in which rare variants explain most heritability (RV model).

### Simulation parameters and fitness functions

In all of the models we considered, our simulations were composed of a diploid population of constant ancestral population size *N_A_* = 500 with three chromosomes, each 500Kb in length. Each chromosome is composed primarily of neutral loci in the first 490Kb, with a locus undergoing selection (described below) in the final 10kb. Of the three chromosomes, only one is undergoing divergent selection (*i.e.*, a change in the phenotypic optimum that corresponds to the trait loci on that chromosome). In general, our simulation parameters were selected to a) qualitatively mirror patterns in the European Corn Borer genome, in which genome-wide *F_ST_* between populations is small but there is high divergence at specific loci (see Results), and b) exemplify extreme ends of the spectrum in terms of genomic architecture, including models in which adaptation is driven primarily by rare alleles or common alleles.

We performed a burn-in of 6*N* generations and verified that this results in steady-state mean trait values in each simulated population, and that a longer burn-in would have a minimal effect on heterozygosity and trait values (Fig. S10). In all simulated models, selection acts on quantitative trait loci (QTL) through a Gaussian fitness function where the strength of selection is determined by *ω*. The fitness effect of stabilizing selection due to the quantitative trait *f_i,j_* of individual *i* with trait value *z_i,j_* and optimal phenotype value *ϕ* is given by

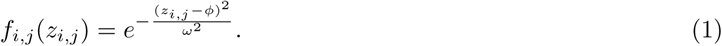

Since we simulate multiple quantitative traits, the composite fitness effect of the *j* traits in individual *i* is given by the product 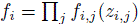.

We considered a range of distinct “genomic architectures” in our simulations, each of which results in a different joint distribution of allele frequencies and effect sizes at equilibrium (after burn-in and before any non-equilibrium changes in the selection model). In the main text, we compare two models that differ widely in their architecture – one in which common variants (CV model) explain most heritability, and another in which rare variants explain most heritability (RV model). Though there are many types of evolutionary models in which rare variants can explain a substantial portion of the heritability, we opted to add an additional directional selection term that penalizes large-effect mutations more than small-effect mutations. Biologically, this represents the possibility of selection acting on variants at multiple scales, for example at the organismal level (through the quantitative trait) and at the molecular level (through gene regulation, protein stability, enzyme kinetics, etc). The directional selection term is written as

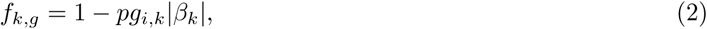

where *k* represents the locus, *|β_k_|* is the absolute value of the effect size of the derived allele at locus *k*, *p* is a constant of proportionality that controls the relationship between selection and effect size, and *g_i,k_* is the number of derived alleles at locus *k* in a given individual. In other words, this is a standard genic directional selection model in which the selection coefficient *s* is represented as *p|β_k_|*, such that alleles with large effects on the trait are also under stronger directional selection. The only difference between the rare-variant model and the common-variant model is that *p* = 0 in the common-variant model and *p* = 2 in the rare-variant model. Both models have exactly the same distribution of effect alleles (see Table 1). Hence, each individual *i* has fitness given by

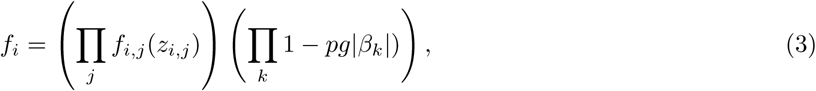

where the right-hand term in the product reduces to 1 in the CV model.

**Table 1:** Simulation parameters used in the CV and RV models. Parameter values corresponding to the CV and RV models are indicated in the corresponding columns. Most RV values are the same as CV values, as indicated by a “-” in the column.

| Parameter | description | CV value | RV value |
| --- | --- | --- | --- |
| $N$ | population size | 500 | - |
| $\mu$ | mutation rate | $2 \times 10^{-7}$ | - |
| $L$ | chromosome length | 500kb | - |
| $c$ | number of chromosomes | 3 | - |
| $r$ | recombination rate | $3 \times 10^{-5}$ | - |
| $L_{\text{QTL}}^a$ | length of QTL locus | 10kb | - |
| $\beta$ | effect size for large-effect mutations | $\pm 0.1$ | - |
| $p$ | scalar that determines relationship between $\beta$ and $s$ | 2 | 0 |
| $q$ | proportion of mutable loci that are effect loci in QTL | 0.9 | - |
| $n$ | number of sampled individuals | 20 | - |
| $B$ | Burn-in, in generations | $3 \times 10^3$ | - |
| $\phi_0$ | initial optimal phenotype value | 1 | - |
| $\delta\phi$ | change in optimum per generation | 0.002 | - |
| $\omega$ | strength of selection | 0.5 | - |
| $m$ | symmetric migration rate after population divergence | varies | - |

<sup>a</sup>There are three selected traits in each simulation, each of which corresponds to a QTL locus on a different chromosome. Each trait has the same initial selection parameters, but the optimal value moves each generation for one of the traits after the Burn-in. Effect mutations are located within a 10-kb locus within each 500-kb chromosome. See Methods section for more information.

In our simulations, at generation 3,000 the population splits into two subpopulations (p1 and p2) of equal size. We allowed symmetric migration between the two subpopulations, ranging in magnitude from *m* = 0.01 to *m* = 0.2, corresponding to high rates of gene flow that might be expected in the early stages of parapatric speciation or situations in which temporal/spatial isolation is incomplete. We picked this range of migration rates to be broadly consistent with genomic patterns and demographic models (using *∂*a*∂*i-cli [Gutenkunst et al., 2009, Huang et al., 2023]) fit to genomic data from our study organism *Ostrinia nubilalis*, as described in section pertaining to analyses of ECB genomic data.

Following the population split, the optimal value *ϕ* corresponding to one of the simulated quantitative traits – which we refer to as the ‘foreground’ – in p2 shifted at a rate of 0.002 per generation, which gradually increases the population mean of the trait value in this subpopulation. We used PIXY [Korunes and Samuk, 2021] to calculate *F_ST_* values over time in each locus throughout the simulation. We continued the simulation for an additional 600 generations (1.2*N* generations) after the population split, sampling the genomes of 20 individuals (40 chromosomes) from each population every 30 generations.

In general, effect size distributions for rare alleles are not known – this is a consequence of the poor statistical performance of association tests for rare alleles. In the absence of any such empirical data to guide our simulations, we opted for a point-mass distribution of effect sizes herein. This allows us to cleanly attribute most genetic variance to rare alleles in the RV model and common variants in the CV model – if variants had broadly variable effect sizes in our simulations, then the differences between the models would be more subtle. While this is useful for interpreting our simulations from a model-based perspective, it is also a limitation for biological interpretation, because it is not likely that genetic variance falls so neatly into these extremes for any real polygenic trait. For the simulations presented in the main text, we used very strong selection (*ω* = 0.5), which is consistent with speciation. We made this choice for two reasons – first, to mimic the biology of the ECB system, which is considered a model for an ongoing speciation event [Dopman et al., 2010], and second, because strong selection is required (but not sufficient) for rare alleles to make substantial contributions to genetic variance. In supplemental analyses, we investigate performance in the context of relaxed selection (see **Results**).

### Likelihood calculation

Our likelihood calculation follows the approach used in sweepFinder [DeGiorgio et al., 2016, Nielsen et al., 2005], but applied to the joint (2D) frequency spectrum as in Nielsen et al. [2009]. The likelihood function is written as

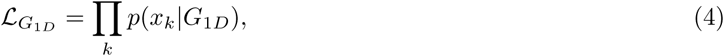

where *p*(*x_k_|G*_1_*_D_*) is the probability of observing a SNP at frequency *x_k_* given the one-dimensional genome-wide frequency spectrum *G*_1_*_D_*. The probabilities are computed using a multinomial, assuming that the height of the frequency spectrum is proportional to the probability of observing a SNP at the corresponding frequency. SweepFinder compares this background *L_G_*_1_*_D_* to *L_F_*_1_*_D_*, which has the same functional form but computes the frequency spectrum based on a set of SNPs found in a given foreground window 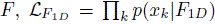. Nielsen *et al* proposed a test statistic *T* that scans for aberrant frequency spectra in local windows,

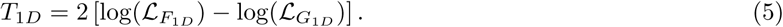

*T*_1_*_D_* is elevated when the likelihood corresponding to the local window is much higher than the likelihood computed using the genome-wide frequencies. Here, we implement a scan using the 2D frequency spectrum, similar to the *G*_2_*_D_* test implemented in Nielsen et al. [2009]. We use the same logic as SweepFinder and use the 2D spectrum to compute *T*_2_*_D_*.

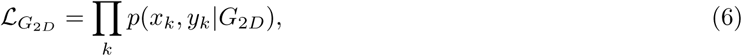

and

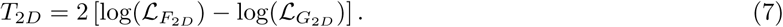

Here, we have replaced the 1D spectrum with the 2D spectrum, taking each bin of the 2D spectrum as representative of the probability of observing a sample from a multinomial. The random variables *x_k_* and *y_k_* represent the frequencies *x* (population 1) and *y* (population 2) of the *k*-th SNP observed in a sample of sequences obtained from the two populations. The allele frequencies can be folded if the ancestral state is not known – in all of our analyses we assume that the ancestral state is unknown and polarize alleles by taking the minor allele in the combined (two population) sample.

*T*_2_*_D_* is sensitive to aberrant frequency spectra, regardless of whether the aberrations are in population 1, population 2, or both populations. We note that *L_G2D_* can be written as

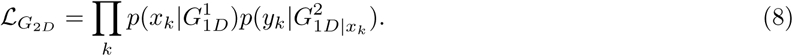

In other words, the probability of observing SNP *k* with joint frequencies (*x*, *y*) is the same as the probability of observing *x* conditional on the 1D-spectrum in population 1, multiplied by the probability of observing *y* in population 2 conditional on the 1D-spectrum in population 2 and *x*. In this form, we can see that taking the log of the likelihood ratio results in

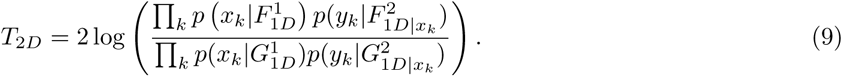

Rearranging terms, we find that

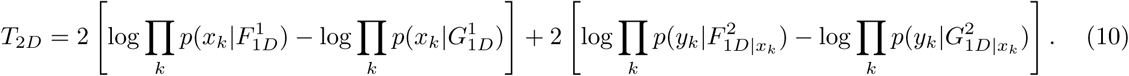

The term on the left is identifiable as *T*_1_*_D_*, while the term on the right is a new term that represents the total contribution of the 2D-spectrum and the 1D-spectrum of the other population. Note that we are also free to decompose *T*_2_*_D_* using the 1D-spectrum of the other population. This demonstrates that *T*_2_*_D_* is likely to be correlated to *T*_1_*_D_* in most circumstances. However, the strength of this correlation depends on the evolutionary history of the populations (which determines the covariance structure of allele frequencies) and how selection shapes the univariate frequency spectra of each population.

### Likelihood-based selection inference

We calculated *T*_2_*_D_* within foreground genomic windows, using the background spectrum as a reference (we discuss the choice of background below). We wrote custom Python scripts to perform these calculations on non-overlapping windows of a fixed size. Our software extracts frequency data from VCF files and a population mapping file that contains information of individual samples and their respective populations. For each genomic window, we computed the folded 1D and 2D SFS using allele frequency data from the sampled populations, where each entry in the folded SFS corresponds to the frequency of the minor allele. For empirical analyses of ECB genomes, the “background” spectrum was computed using the whole genome. For analyses of simulated populations, we compared foreground windows to two backgrounds, one of which was neutral and another of which had only stabilizing selection. See the **Power analyses** section for more information about background spectra and comparisons between the foreground windows and genomic background. In general, the choice of background can have a substantial impact on the performance and interpretation of scans based on *T*_1_*_D_* and *T*_2_*_D_*, as is true for other types of genomic scans – we discuss the impact of the choice of the background in the results section.

There are multiple options for identifying candidate regions under selection within our framework, falling into two broad categories – approaches that use a parametric model-based null, and non-parametric approaches that simply scan for outliers. In this study, we opt for an outlier scan because we are primarily interested in developing an approach that is very straightforward to apply to non-model genomes. As discussed in Nielsen et al. [2005], the null distribution of *T*_1_*_D_* (and consequently also *T*_2_*_D_*) depends on evolutionary processes such as recombination and population demography. Therefore we identify outliers simply as the upper tail of the distribution of the test statistic of interest, noting that the approach could also be used with a model-based null distribution, which could be obtained through simulations or analytical calculations. We used SciPy to calculate multinomial probabilities [Virtanen et al., 2020]. Our scripts are freely available at https://github.com/alecalderonb/2DSFS-scan.

#### Power analyses

A major question addressed in this study concerns the statistical performance of *F_ST_* scans relative to likelihood-based scans for different genomic architectures and evolutionary models. To this end, we calculated statistical power at different significance thresholds for both *F_ST_* and *T*_2_*_D_*.

We define power at a specific significance threshold *α* as the probability that a genomic window under divergent selection has a test statistic value that falls in the right-most (1-*α*) tail of the distribution of test statistics computed from the genome background. We compute the distribution of background test statistics using two distinct sets of background loci. In the main text, we compare test statistics at foreground loci to background loci that have the same trait model as the loci under divergent selection, but no divergent selection (*i.e.*, no change in optimal trait value *ϕ* after the population split). In supplemental analyses, we also compare the foreground loci to a neutral locus with no selection. The former analysis is more conservative and more informative about divergent selection – when we compare to a neutral locus, any selection (whether divergent or simply due to constraint) will distort the frequency spectrum and result in elevated test statistics. This more conservative analysis is analogous to comparing missense sites in a gene or group of genes to other missense sites in the genome background, rather than comparing missense sites to noncoding sites. Such missense-to-missense comparisons are commonly performed in selection analyses (see for example Hernandez et al. 2011).

For all power analyses, we performed 1,000 independent simulations in SLiM of a given set of parameters. Each simulation produced a VCF file as output, and we combined all 1,000 VCFs into a single file. This allowed us to make a composite genome that is comparable in size to the number of unlinked sites (*≈* 2.2 *×* 10^6^) in the our European Corn Borer analyses (see section on **Analysis of ECB genomic data**). We then computed both *T*_2_*_D_* and *F_ST_* in windows of 500 SNPs. We used PIXY [Korunes and Samuk, 2021] to compute *F_ST_* and custom Python software to compute *T*_2_*_D_*. Because the sites in the analyses represent 1,000 independent simulation runs, LD between simulated polymorphic sites is weak.

In general, our power analyses should only be taken as comparative between the specific evolutionary models that we simulate, and cannot be generalized to all rare– or common-variant architectures. Unlike single locus statistical tests, it is not straightforward to query statistical performance across the full space of model parameters, because every evolutionary model (and the specific parameters corresponding to that model) will generate a distinct joint distribution of allele frequencies and effect sizes. Moreover, effect mutations in our simulations are concentrated within an effect locus – more diffuse genetic architectures would be more difficult to capture with this approach. Consequently, our simulated models are only intended to represent extremes of genetic architecture, and should not be taken as practical recommendations for specific use cases. We review potential implications and limitations of our specific choices in the Discussion.

### Analysis of *Ostrinia nubilalis* (ECB) genomic data

To assess patterns of genetic variation and differentiation between ECB populations, we used individual-level sequence data from populations that differ in voltinism (the number of generations per year) a trait that previous studies have found to generate reproductive isolation and be under divergent selection [Dopman et al., 2010, Kozak et al., 2019, Kunerth et al., 2022]. Sequence data were obtained from the National Center for Biotechnology Information Sequence Read Archive (BioProject ID: PRJNA540833), including 18 univoltine (uv) and 14 bivoltine (bv) individuals collected from field sites across New York State, as described in Kozak et al. [2019]. Since we are interested in the genomic basis of reproductive isolation between these phenotypically distinct groups – which are isolated temporally due to differences in the timing of seasonal emergence – we analyzed them by pooling individuals by their voltinism phenotype, except where otherwise noted. ECB populations are also behaviorally isolated due to genetically encoded sex pheromone [Lassance et al., 2010, Unbehend et al., 2021], and all individuals were of Z pheromone type. We chose a subset of individuals that corresponded to geographically close subpopulations and for which we had high-quality phenotype data available.

Raw sequencing reads were processed using fastp [Chen et al., 2018] to remove adapter sequences and filter reads shorter than 150 base pairs. High-quality reads were then aligned to the *O. nubilalis* reference genome (ilOstNubi1.1, [Boyes et al., 2025]) using the Burrows-Wheeler Aligner (BWA) [Li and Durbin, 2010], with secondary hits marked. To improve data quality, we removed low-quality alignments and PCR duplicates using Picard Toolkit [Pic, 2019]. The resulting BAM files were indexed with Samtools [Li et al., 2009].

Single nucleotide polymorphisms (SNPs) were called using bcftools [Danecek et al., 2021] and samples were grouped per geographic location using the-G option so that the Hardy-Weinberg Equilibrium assumption was applied within but not across populations. We applied filtering criteria using vcftools [Danecek et al., 2011] to retain high-confidence variants, removing SNPs with a depth below 10 or above 100, a minimum quality score of 40, and a call rate tolerance of 80%. Our variant calling pipeline applied to 18 univoltine and 14 bivoltine individuals resulted in 9,093,357 SNPs. We used PIXY [Korunes and Samuk, 2021] to calculate *F_ST_* in windows of various sizes.

We used *∂*a*∂*i-cli [Gutenkunst et al., 2009] to infer the demographic parameters that were used in our simulations. We used the observed 2D SFS extracted from biallelic synonymous SNPs to fit a split-with-migration model. In this model, an ancestral population diverges into two subpopulations that undergo an instantaneous change in size, followed by continuous migration between them. We selected the best-fitting population size and symmetric migration rate parameters based on likelihood-based model comparisons.

In an effort to limit our vulnerability to false positive selection signals driven by linkage disequilibrium [Bustamante et al., 2001], we used plink 1.9 [Purcell et al., 2007] to LD-prune ECB genomic data for analyses using both *F_ST_* and *T*_2_*_D_*. For genome-wide analyses presented in the main text, we LD-pruned to an *r*^2^ of 0.5 in 10kb windows, resulting in 2,163,452 retained SNPs. We annotated synonymous and nonsynonymous sites using SnpEff with a custom database made with the ECB reference genome [Cingolani et al., 2012], resulting in 423,355 retained SNPs. LD-pruning using similar methods (*r*^2^=0.5) on the coding regions further reduced this number to 192,155 retained SNPs, and we estimated a mean per-site missingness in this dataset of 0.14%, and per-individual missingness ranging from 0.02% to 1.05%. We use these two sets of SNPs (genome-wide and coding) for analyses presented in the main text, and consider downstream implications of LD-pruning in the Discussion section.

In insects, prior work suggests that circannual rhythms are linked to the daily circadian rhythms and genetic control of both traits maps to the circadian clock pathway [Helfrich-Förster, 2024, Meuti and Denlinger, 2013]. In ECB, daily circadian rhythms differ among uv and bv individuals, with knockout of the *period* gene leading to loss of daily circadian rhythms, in addition to influencing seasonal timing [Dayton et al., 2025, Kozak et al., 2019]. We used KEGG Release 117.0+/02-06 to find genomic coordinates of loci containing genes with known functions related to circadian or circannual rhythms [Kanehisa et al., 2025] in ECB. Genes that are members of the circadian circuit (pathway onu04711 in KEGG) we identified as “core” genes (Tab. 2), while genes implicated in circadian cycles that were identified in other empirical studies (largely performed in *Drosophila melanogaster*) are included as “non-core” genes [Grima et al., 2012, Kula-Eversole et al., 2021, Petsakou et al., 2015, Sekiguchi et al., 2024, Wang et al., 2020].

**Table 2:**
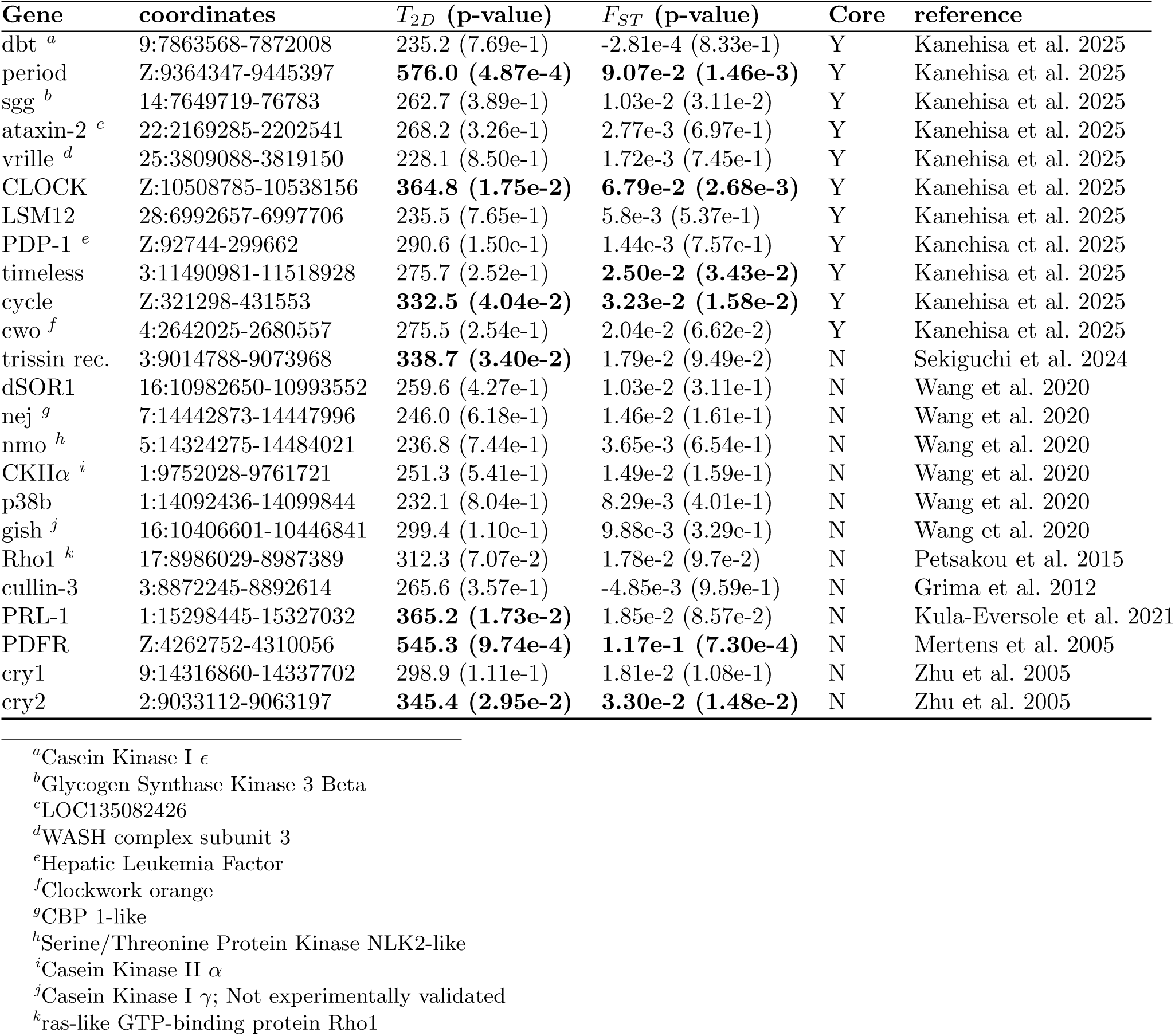
Test statistics for circadian clock genes for windows of 500 SNPs using the whole genome (not only coding loci). Gene names in the **Gene** column correspond to naming conventions in *Drosophila melanogaster*, which sometimes differ from names in the annotated ECB genome. Names that correspond to the ECB genome build (ilOstNubi1.1, December 2023, Boyes et al. 2025) are given in footnotes. **Core** genes are members of the ECB circadian pathway (pathway onu04711) in KEGG release 117.0 [Kanehisa et al., 2025]. The remaining genes have been implicated in other experimental studies but are not part of the core circuit in KEGG. Test statistic values for *T*_2_*_D_* and *F_ST_* correspond to a screen of the whole genome in windows with a fixed width of 500 SNPs, after LD pruning (see Methods). Test statistics that correspond to p-values under 0.05 are written in bold font.

## Results

### *F_ST_* and frequency spectrum patterns for RV/CV models

We first explored how the RV and CV models affect patterns of variation at trait loci over time. Under the RV model, the mean phenotype increases in the population under divergent selection, tracking the change in the optimal phenotype value but with a slight lag (Fig. 2A). *F_ST_* increases over time (Fig. 2B) in both the genome background (gold lines) and at trait loci (blue, background trait; pink, foreground trait). However, differences in *F_ST_* between the foreground and the genome background are subtle until about *t* = 300, and differences between trait-foreground and trait-background loci remain fairly small until late in the time-course. This is contrast to the CV model, in which there was very little lag and somewhat larger *F_ST_* differences were observed (Fig. S1A-B), though differences between background and foreground were still small before *t* = 300. This suggests that *F_ST_* is likely to be powerful at late times in both the CV and RV models, but may lack sensitivity at early time points. This is consistent with our expectation – when rare alleles drive phenotypic differentiation, *F_ST_* should be relatively insensitive. These overall trends held true in both models for a wide range of migration rates, though the magnitude of differences in *F_ST_* between foreground and neutral/background loci varied considerably across migration rates (Fig. S2-S3).

**Figure 2:**
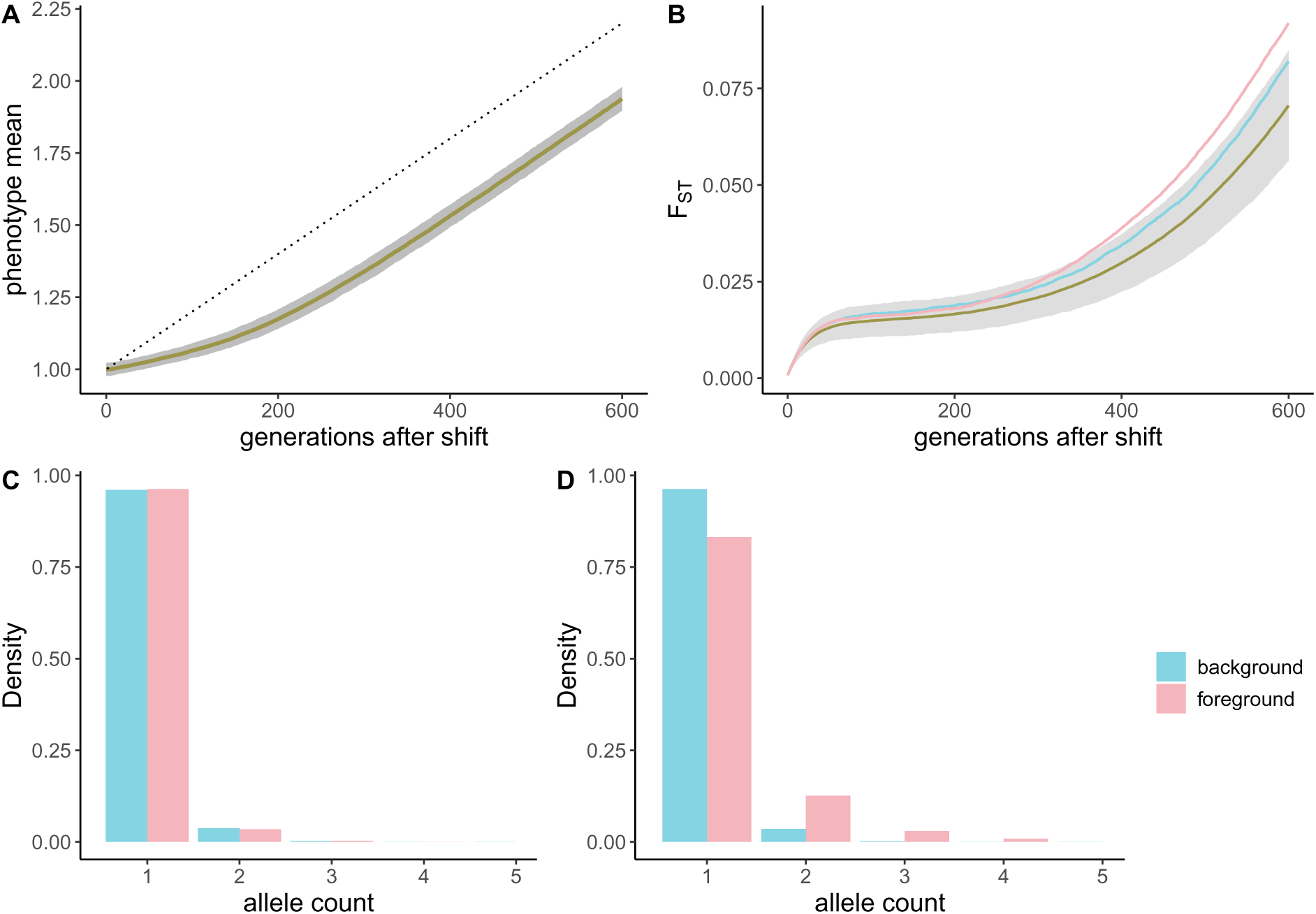
Simulation results for the RV selection model. **A**) The mean of the phenotype under divergent selection in p2 after the population split. The optimal phenotype value is shown with the dotted line. The observed mean is shown in gold across 10^3^ simulation replicates, and the gray represents three standard deviations around the mean. **B**) *F_ST_* at neutral (gold), divergent (pink), and stabilized (blue) loci. The gray envelope represents two standard deviations around the mean for neutral loci. **C**) Frequency spectra for QTLs at the population split (*t* = 0) and **D**) *t* = 300. Blue bars represent the stabilized locus and pink bars represent the locus under divergent selection. Only the first 5 bins of the spectrum (in a sample of 40 chromosomes) are plotted because selection is strong and there are very few alleles in the high frequency bins at these time points.

In contrast, the one-dimensional frequency spectra of trait alleles in population p2 were strongly informative of divergent selection in the RV model by *t* = 300, with a strong observed shift to higher frequencies in the foreground loci (Fig. 2C-D). The one-dimensional frequency spectra in the CV model showed no discernible shift (Fig. S1C-D), suggesting that *F_ST_* differences were being driven by a small number of alleles with larger than average frequency differences rather than a population-specific shift in the shape of the spectrum.

### Comparison of *F_ST_* and *T*_2_*_D_*

Next, we explored the relationship between *T*_2_*_D_* and *F_ST_* in our simulations by computing both statistics in windows with a fixed number of SNPs (500 SNPs; Fig. 3). We observe that *T*_2_*_D_* and *F_ST_* are modestly correlated at early time points (*ρ* = 0.255, *p <* 1*e −* 10; Fig. 3A) in the RV model, while they are highly correlated at late time points (*ρ* = 0.760, *p <* 1*e −* 10; Fig.3B). In the CV model, the two statistics were weakly correlated at both time points, although the correlation is stronger at *t* = 600 (Fig. S4). We then computed power at the *α* = 5 *×* 10*^−^*^3^ threshold based on the distribution of test statistics at background selected loci (see Methods), finding that in the RV model, *T*_2_*_D_* is informative about divergence at early time points, while *F_ST_* is not (Fig. 4). At the later time point, both statistics are informative. Trends in power as a function of time were similar across migration rates for *T*_2_*_D_*, while *F_ST_* was most powerful at intermediate migration rates. This is in contrast to the CV model, in which neither statistic is informative at early time points and only *F_ST_* is informative at *t* = 600 for all non-zero values of migration rate (Fig. S6).

**Figure 3:**
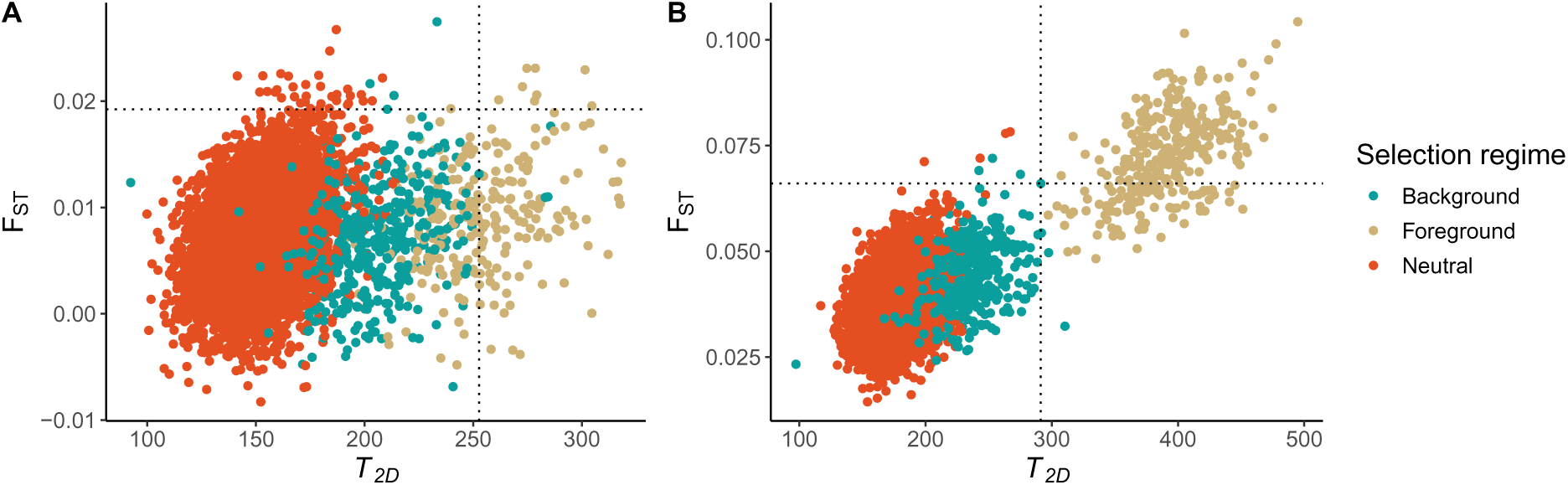
Comparison of *F_ST_* and *T*_2_*_D_* values at *t* = 300. (**A**) and *t* = 600 (**B**) in the RV model. Note that the difference is scale on both axes between the two timepoints, as both *T*_2_*_D_* and *F_ST_* increase over time in the simulation. The dashed lines represent the *α* = 5 *×* 10*^−^*^3^ threshold, as computed using the background selected loci.

**Figure 4:**
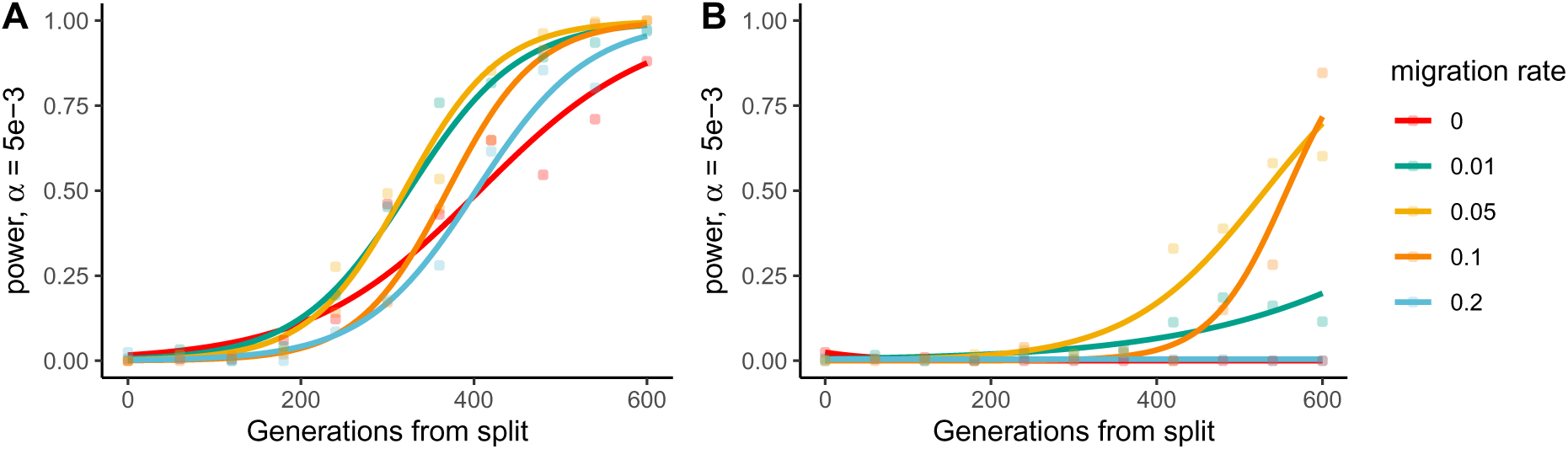
Power at *α* = 5 *×* 10*^−^*^3^ for the RV model for *T*_2_*_D_*. (**A**) and *F_ST_* (**B**). Each point represents the fraction of loci undergoing divergent selection that had test statistics in the (1-*α*) right tail of the background test statistic distribution. Colors correspond to various migration rates, and the solid lines represent a binomial regression on the data.

We also sought to explore how power would depend on the strength of selection, especially since we used very strong selection for most simulations in the main text. In Fig. S12, we plot power as a function of time for several values of *ω*. In these simulations, we set all model parameters to the same values we used for the RV model, except for the rate of change of the optimum phenotype (*δϕ*), which we scale proportional to the strength of selection (*i.e.*, the square of *ω*). This additional parameter modification is required, because without it little adaptation occurs at weaker selection values on these timescales since trait variance is large relative to the change in the optimum (see Fig. S11). We observed that *F_ST_* performs better at earlier times as selection strength decreases, while *T*_2_*_D_* performs worse at later times for weak selection. This is consistent with our expectation, since weaker selection results in architectures that are dominated by common alleles.

When comparing foreground loci to a background composed of neutral loci, *T*_2_*_D_* is informative at all times (Fig. S6) – however, this is not a fair comparison, because the 1D spectrum is shifted to much lower frequencies for sites under stabilizing selection – *T*_2_*_D_* would be informative in this comparison even if there were no divergence due to the effects of stabilizing selection on the frequency spectrum. We include this analysis only for completeness. We also estimated ROC curves for three time points (*t* = 0, 300, 600) using the *T*_2_*_D_* data, which showed that *T*_2_*_D_* has good overall performance in distinguishing between neutral and diverged windows in the RV model (Fig. S9).

### Empirical analysis of ECB populations

We investigated empirical patterns of variation captured by *T*_2_*_D_* and *F_ST_* in European Corn Borer genomes corresponding to the uv (univoltine) and bv (bivoltine) ecotypes. These ecotypes are undergoing partial reproductive isolation due to differences in seasonal timing of emergence [Kozak et al., 2019]. Prior *F_ST_* scans identified a genomic regions that are highly differentiated between uv and bv populations on the Z chromosome [Kozak et al., 2019, Kunerth et al., 2022], corresponding to the loci containing the *period* and *pigment-dispersing factor receptor* (*pdfr*) genes, both of which are part of circadian rhythm pathways (Tab. 2). We compared folded 1D and 2D frequency spectra within the *pdfr* locus to the genome background to explore how allele frequency patterns differ between diverged and well-mixed genomic regions (Fig. 5). Though 1D spectra differ slightly in shape between the foreground and background regions in both the univoltine and bivoltine populations (Fig. 5A-B), comparison of the 2D spectra reveals that differentiation between populations is much higher in the region of interest on the Z chromosome (Fig. 5C-D). This suggests that both *T*_2_*_D_* and *F_ST_* are likely to be elevated in this region. Indeed, an *F_ST_* scan of these genome results in patterns that are concordant with the aforementioned prior studies, with large peaks on the Z chromosome corresponding to the loci containing *pdfr* and *period*.

**Figure 5:**
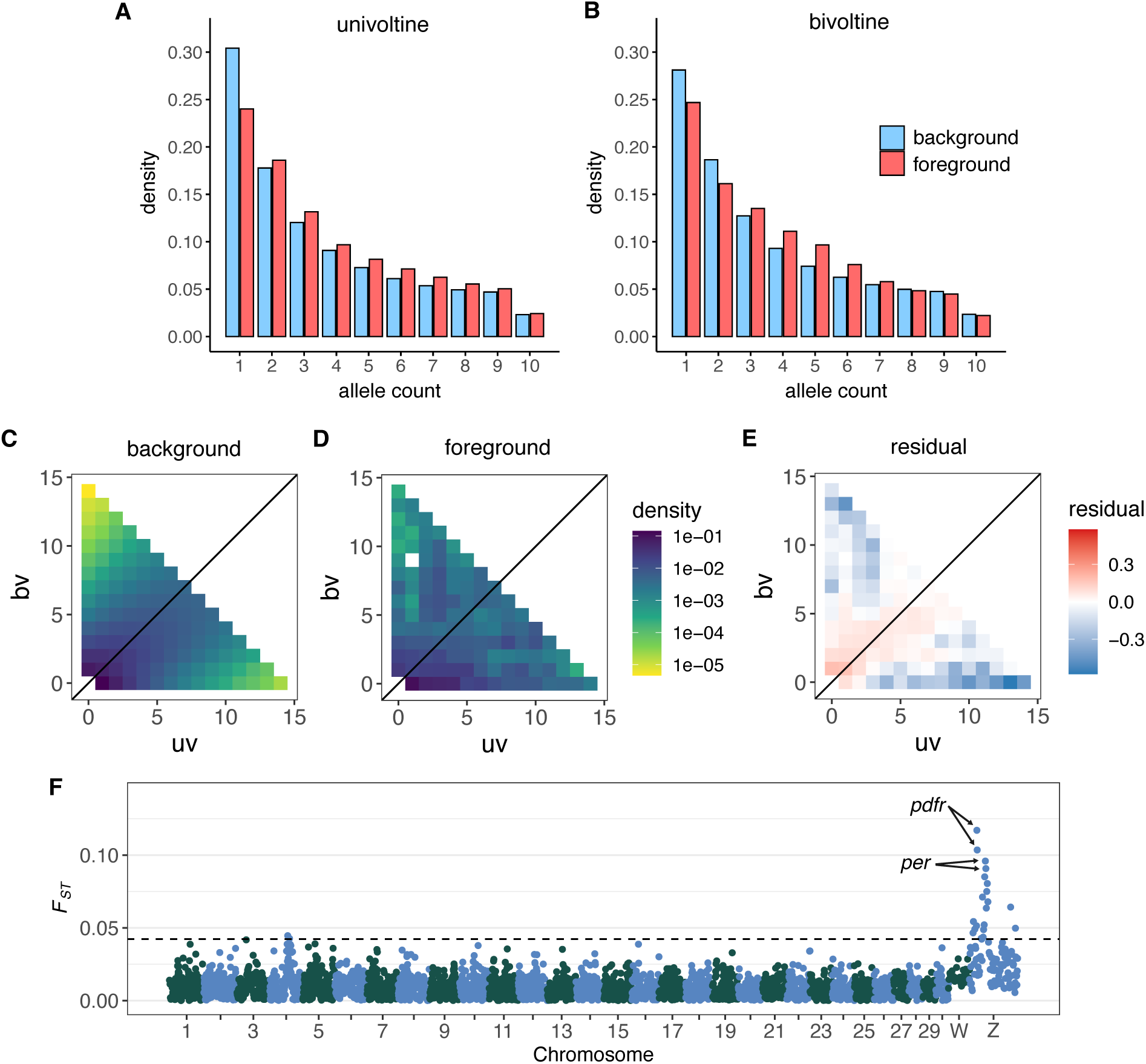
Patterns of genetic variation in the ECB genome. (**A**) Frequency spectra for alleles at two genomic regions: background (autosome 1, blue bars) and foreground (500 kb window in chromosome Z, red bars) in the univoltine (uv) and (**B**) bivoltine (bv) individuals. (**C**) Two-dimensional frequency spectrum at the background and **D**) foreground region. **E**) Poisson residuals between the background and the foreground 2D spectra. **F**) *F_ST_* scan in windows of 500 SNPs across the ECB genome.

We sought to compare empirical patterns in *F_ST_* to *T*_2_*_D_* across the ECB genome, supposing that some loci that are sensitive to *T*_2_*_D_* might have been missed with *F_ST_*. We LD-pruned the genome for these analyses to limit our false-positive rate due to linked sites. In bins of varying width (from 250 to 1,000 SNPs) we computed both *F_ST_* and *T*_2_*_D_*. Similar to our findings in simulated data at later time points with the CV model, we observed that *T*_2_*_D_* and *F_ST_* were correlated, especially on the Z-chromosome (Fig. S7; Tab. S1). However, some loci with elevated *T*_2_*_D_* values do not have elevated *F_ST_* – while this does not guarantee that loci discovered with *T*_2_*_D_* alone are not spurious, it does suggest that the two statistics can provide different sets of putatively positive loci. This discordance is consistent with *T*_2_*_D_* capturing selection signals not detected by *F_ST_*, though LD-pruning alone might not ensure that the false-positive rate of *T*_2_*_D_* at these loci matches that of *F_ST_*.

In general, sites with stronger evolutionary conservation should have elevated *T*_2_*_D_* values relative to the genome background. We therefore further limited our analysis to include only coding loci (see Methods), and repeated our scan in windows of 250 SNPs. We reduced the window size to 250 because by subsetting the genome to include only non/synonymous sites we necessarily increase the physical size of windows with a fixed number of SNPs. We found that *T*_2_*_D_* and *F_ST_* are both elevated in previously identified loci on the Z chromosome corresponding to *pdfr* and *period* (Fig. 6). Several additional loci are identified with elevated values of *T*_2_*_D_*, but not *F_ST_*. Two of these regions contain previously identified circadian genes, *rho1* (chromosome 17) and *prl-1 phosophatase* (chromosome 1; Kula-Eversole et al. 2021, Petsakou et al. 2015). A third region on chromosome 3 with an even higher value of *T*_2_*_D_* contained several genes, but none with immediately obvious circadian functions. We then sought to characterize which bins of the frequency spectrum contributed to the high values of *T*_2_*_D_* in these two loci. We found that a slight excess of population-specific low frequency SNPs (*i.e.*, variants with frequencies that were substantially higher in one population than the other but not high frequency in the pooled sample) contributed to *T*_2_*_D_* in both loci, but higher frequency bins also contributed to the signal (Fig. S8).

**Figure 6:**
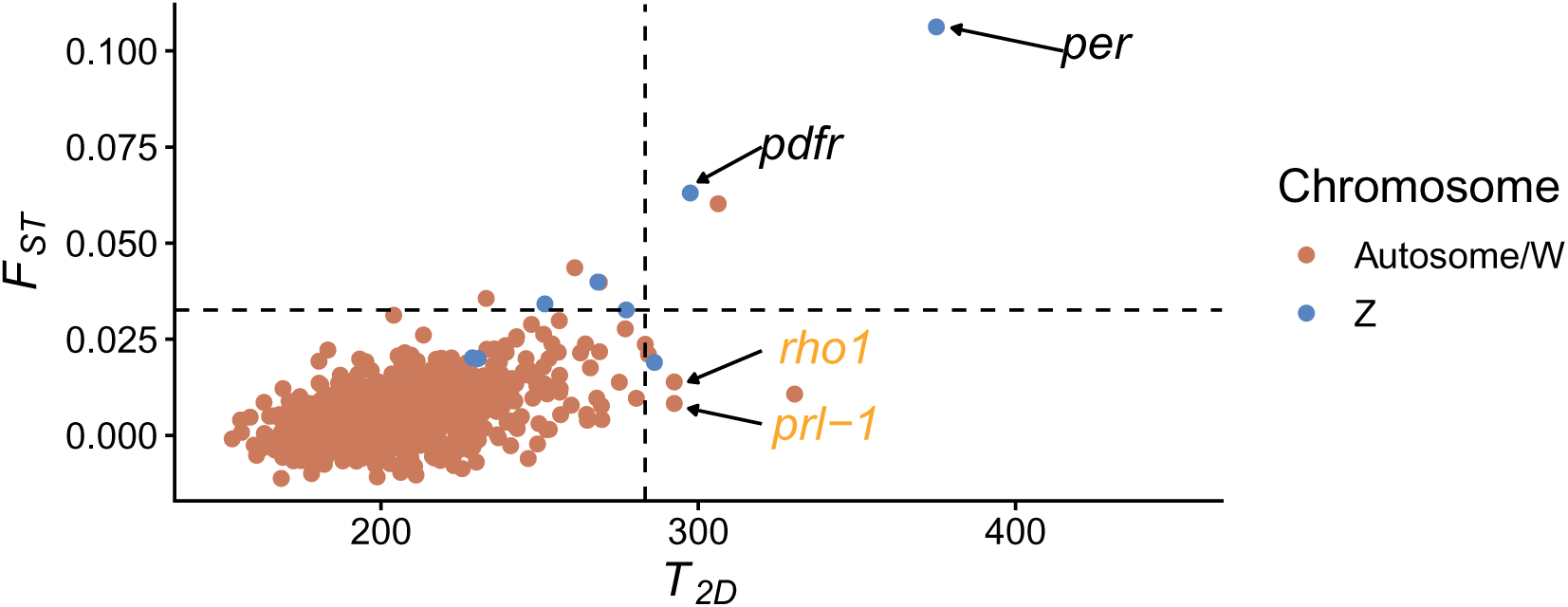
Comparison of *T*_2_*_D_* and *F_ST_* values calculated using only nonsynonymous and synonymous sites in ECB genomic data in windows of 250 SNPs. The dashed line represents the *α* = 1*e −* 2 threshold. Loci on the autosomes or W chromosome are shown in salmon, while loci on Z are shown in blue. The names of genes in the ECB circadian pathway that fall within each locus are indicated in black text, while genes that have been implicated in circadian rhythms are indicated in orange text.

We also computed *T*_2_*_D_* and *F_ST_* in windows of 500 SNPs to scan the whole genome for elevated selection signals at circadian genes that were identified in prior studies [Grima et al., 2012, Kula-Eversole et al., 2021, Petsakou et al., 2015, Sekiguchi et al., 2024, Wang et al., 2020] or included in the circadian pathway for ECB as reported in KEGG (see Methods; [Kanehisa et al., 2025]) (Tab. 2). In addition to *period* and *pdfr*, four other genes were identified as outliers at the *α* = 5 *×* 10*^−^*^2^ level with *F_ST_* and five were identified with *T*_2_*_D_*. Of the two detected with *T*_2_*_D_* alone, one (*prl-1*) was also identified in our scan of the coding portion of the genome while the other (*trissin* receptor) was not. The locus containing *rho1* had a somewhat elevated value of *T*_2_*_D_* in this latter genome scan, but missed the cutoff for significance.

## Discussion

We evaluated the utility of a likelihood-based framework for detecting genomic regions of differentiation, particularly at early stages of the differentiation process in a model in which rare variants drive trait divergence. By comparing the joint frequency spectrum in foreground windows to the background genome, we showed that a likelihood-based statistic (*T*_2_*_D_*) can identify divergent loci even in the presence of high gene flow when rare alleles drive trait divergence, while *F_ST_* has low power to detect such regions. When common variants are the primary drivers of phenotypic divergence, the two approaches performed similarly. These simulation-based results were reflected in an application to empirical data, where two previously detected loci were identified by both statistics, while several new candidate loci were identified by *T*_2_*_D_* alone. Our results demonstrate the potential utility of likelihood-based approaches for detecting previously-undiscovered loci undergoing divergent selection.

Our findings are consistent with theoretical expectations about the early stages of adaptation between populations undergoing partial reproductive isolation. When the optimal value of a selected trait changes in a population, the early stages of phenotypic differentiation can proceed rapidly through alleles of large effect, even while those alleles remain relatively rare in the population [Hayward and Sella, 2022, Stetter et al., 2018, Uricchio et al., 2019]. Such variants can produce substantial shifts in trait means even before these alleles become common, because of their large effect sizes. As a result, early adaptation may be driven in part by alleles that contribute disproportionately to phenotypic change but leave only subtle signatures in traditional population summary statistics. In particular, such alleles will rarely fix, because once the trait mean approaches the new optimal value selection acts against these variants [Hayward and Sella, 2022]. Statistical methods that directly leverage the full distribution of allele frequencies are therefore promising avenues for the detection of these early signals of differentiation. As we discuss below, several practical challenges would need to be overcome to improve the interpretability of such approaches.

Our analysis revealed additional loci of interest for potential follow-up in studies of differentiation between univoltine and bivoltine European Corn Borer populations. *T*_2_*_D_* and *F_ST_* both identified divergence in the circadian clock genes *cycle*, *clock*, and *cryptochrome2*, all three of which have been found to contribute to regulation of temperature-induced diapause in *Bombyx* moths [Homma et al., 2022]. Two of the top three loci that were *T*_2_*_D_* outliers but not *F_ST_* outliers contained genes previously identified as members of the circadian circuitry in *Drosophila melanogaster*. *Rho1* was implicated in rhythmic control of neural plasticity and seasonal adaptation [Petsakou et al., 2015], while *prl-1* dependent dephosphorylation of *timeless* in neurons sets the period length and phase of the circadian clock in low light conditions [Kula-Eversole et al., 2021], and potential epistatic effects on *Pdfr* have been suggested [Li et al., 2022]. These results point to a potential role for adaptation in the regulatory elements that modulate the circadian pathway and contribute to further reproductive isolation between uv and bv populations, perhaps subsequent to adaptation in the core genes themselves.

Although we believe that there is substantial under-utilization of likelihood-based approaches in the domain space of detecting early local adaptation, it must also be recognized that there are substantial remaining challenges that could limit their broad utility if unaddressed. Three of the most important include the diffuse signal of heritability across the genome for many heritable traits, linkage and genome structure, and confounding with signals of selective constraint. We discuss each of these issues in turn and consider possible remedies.

For many complex heritable traits, heritability is diffusely spread across the genome, and most associations with traits are weak and explain a small proportion of trait variance [Boyle et al., 2017]. This is potentially problematic for the application of likelihood-based methods if our goal is to detect loci with rare alleles underlying phenotypes related to reproductive isolation – if these alleles are equally as broadly distributed as common allele associations, there is little hope to discover them with such an approach, because their impact on the allele frequency spectrum will be very subtle. Indeed, our simulations assumed that a single 10kb locus contained a substantial mutation rate for trait altering-alleles, a situation that may not always match biological reality. However, there is reason to believe that rare, large-effect alleles may not be as broadly distributed as GWAS hits. First, rare variant association tests have been widely deployed to detect loci harboring causal rare alleles [Lee et al., 2012, Neale et al., 2011], and rare alleles appear to play important roles in many phenotypes, including RNA expression variation, heart disease, human height, and prostrate cancer [Chen et al., 2022, Do et al., 2015, Frésard et al., 2019, Mancuso et al., 2016]. From an evolutionary perspective, it is likely that alleles with large effects will occur in genes or regulatory loci that are central (rather than peripheral) to the phenotype of interest, and consequently may be more likely to fall into a small number of loci than common alleles with trait associations. In cases where core genes of interest for a trait relevant to population divergence are known (such as the circadian clock circuit considered in this paper), it is possible to combine signals across these genes to assess these loci as a group and boost power. Still, our simulations represented a “best-case scenario” in which rare variants were highly clustered. In practice, causal loci are likely to harbor alleles with a range of effect sizes and frequencies, and many such loci will contain too few loci to be detected with likelihood-based approaches.

Linkage across the genome presents a well-known challenge for likelihood-based approaches that utilize the frequency spectrum. Correlations between frequencies are problematic because frequency spectrum-based approaches typically assume that each site represents an independent observation, and neutrally evolving loci are more likely to be detected as false positives when recombination rates are low [Bustamante et al., 2001]. A common solution to this problem is to reduce LD in the data by LD-pruning, as we did here. Though this approach should work to reduce false positives, a challenge is that it is likely to reduce power in unanticipated ways, because alleles are greedily removed from the dataset until no loci have correlations in excess of a user-defined threshold. LD-pruning is also highly frequency-dependent – the correlation coefficient *r*^2^ is bounded depending on the frequencies of a pair of alleles, and the highest potential values of *r*^2^ occur when the frequencies are the same. Ideally, likelihood-based approaches would be able to make use of the full data, but this would require explicit modeling of the linkage between sites or at least accounting for potential variation in recombination rates across the genome, which may be challenging to model. It is also worth noting that the performance of selection statistics can be affected by missing data, especially in low-coverage samples, and this has been studied in frequency-spectrum-based statistics [Pandey et al., 2024], as well as PCA-based statistics [Meisner et al., 2021], though future studies need to be conducted to better understand its effects on *T*_2_*_D_*.

Evolutionary conservation also poses a challenge for the approach we considered in this study. Since we only scan for aberrant frequency spectra, frequencies that skew lower due to selective constraint without any signal of differentiation would also be detected. For this reason, we sought to compare coding loci to other coding loci in our first set of empirical analyses. In principle, signals of constraint could be accounted for by computing *T*_2_*_D_* for specific bins of the spectrum that are most informative for divergent selection, or potentially by computing *T*_2_*_D_* conditional on *T*_1_*_D_* in the combined sample (since the 1D spectrum should also be informative about conservation). Another more sophisticated approach would be to use a model-based background rather than an empirical background, as is possible in sweepFinder [DeGiorgio et al., 2016, Nielsen et al., 2005]. This would require the development of predicted frequency-spectra for divergent polygenic traits under selection, which to our knowledge is only possible in simulation for complex demographies.

Our work suggests that likelihood-based scans could be a promising approach for the elucidation of differentiation in the presence of high gene flow. In such situations, some loci may not be detected with *F_ST_*, especially if the primary drivers of adaptation have low frequencies in the joint sample. We suggest that such approaches could be a fruitful avenue of study and further methods development, and could enrich our understanding of the evolutionary mechanisms underlying reproductive isolation.

## Data availability

Summary statistics computed in this study are freely and publicly available in a GitHub repository at https://github.com/alecalderonb/2DSFS-scan. All primary data were derived from publicly available datasets as described in the main text.

## Acknowledgments

We thank Erik Dopman and members of the Dopman and Uricchio labs for many helpful conversations throughout the development of this project. We are grateful to our anonymous reviewers and Dr. Sam Yeaman, each of whose comments helped us to substantially improve the manuscript.

## Supplementary Data

**Figure S1:**
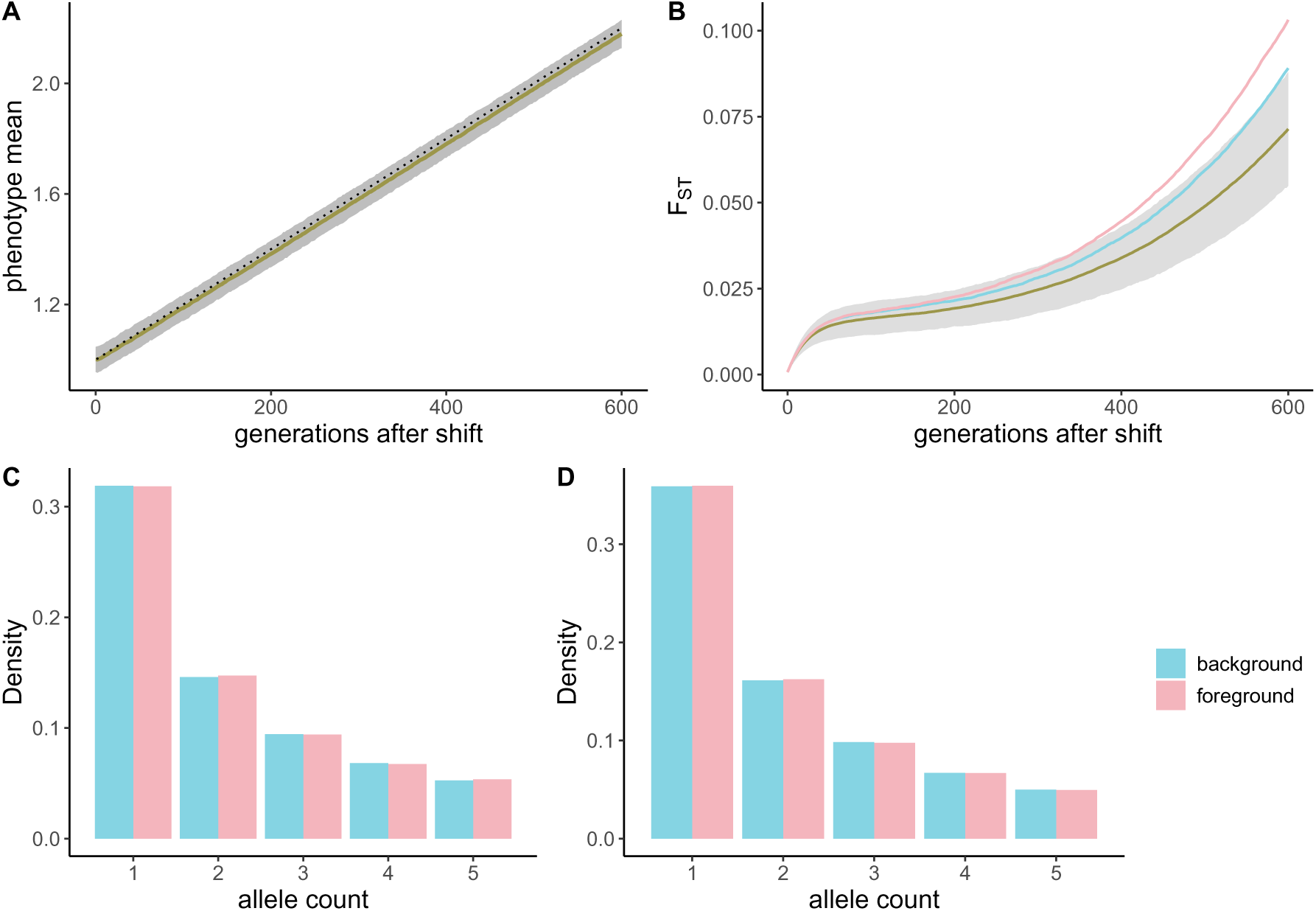
Simulation results for the CV selection model. **A**) The mean of the phenotype under divergent selection in p2 after the population split. The optimal phenotype value is shown with the dotted line. The observed mean is shown in gold across 10^3^ simulation replicates, and the gray represents three standard deviations around the mean. **B**) *F_ST_* at neutral (gold), divergent (pink), and stabilized (blue) loci. The gray envelope represents two standard deviations around the mean for neutral loci. **C**) Frequency spectra for QTLs at the population split (*t* = 0) and **D**) *t* = 300. Blue bars represent the stabilized locus and pink bars represent the locus under divergent selection. Only the first 5 bins of the spectrum (in a sample of 40 chromosomes) are plotted.

**Figure S2:**
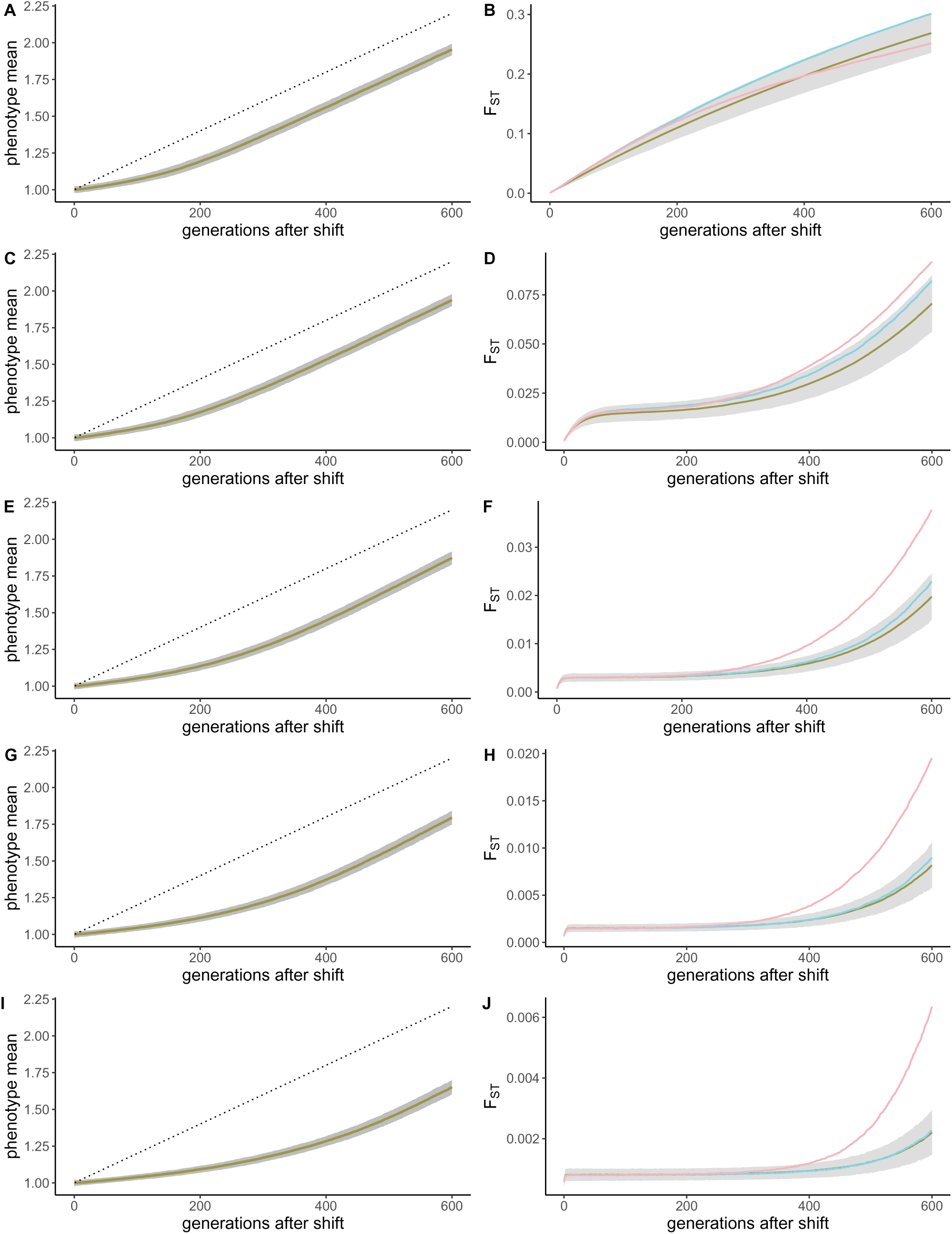
Phenotype mean in p2 and *F_ST_* between p1 and p2 for five migration rates in the RV model. The gold lines in the left hand column represent the phenotype mean over 1,000 independent simulations while the gray envelope represents two standard deviations around the mean. The dashed line represents the optimal value of the phenotype. In the right hand column, the gold line represents the mean *F_ST_* at neutral loci, while the gray envelope represents three standard deviations around this mean. The blue line represents the mean *F_ST_* and background (non-divergent) trait loci, while the pink line represents the mean *F_ST_* in loci that underly trait divergence. Migration rates are *m* = 0 (A-B), *m* = 0.01 (C-D), *m* = 0.05 (E-F), *m* = 0.1 (G-H), *m* = 0.2 (I-J).

**Figure S3:**
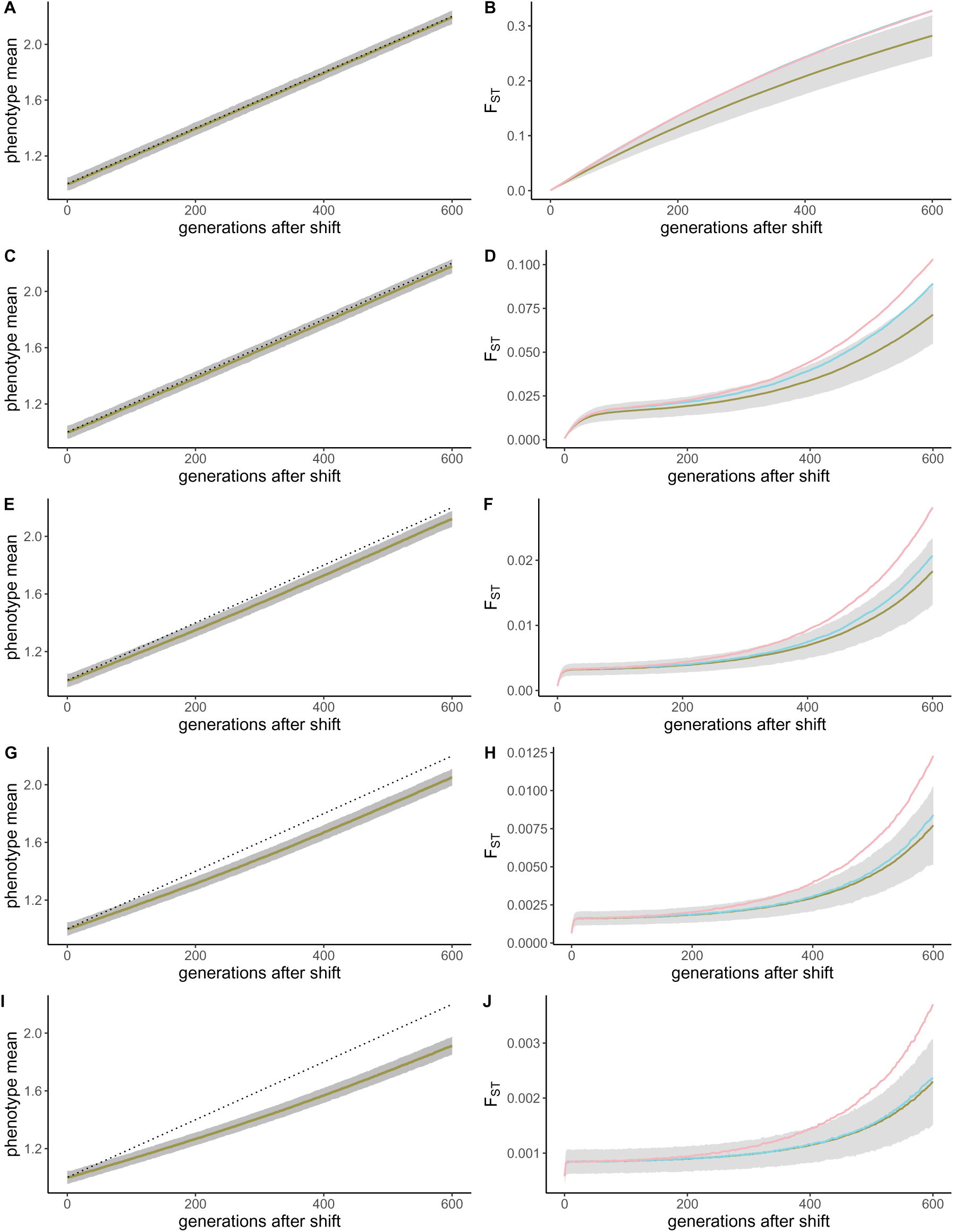
Phenotype mean in p2 and *F_ST_* between p1 and p2 for five migration rates in the CV model. The gold lines in the left hand column represent the phenotype mean over 1,000 independent simulations while the gray envelope represents two standard deviations around the mean. The dashed line represents the optimal value of the phenotype. In the right hand column, the gold line represents the mean *F_ST_* at neutral loci, while the gray envelope represents three standard deviations around this mean. The blue line represents the mean *F_ST_* and background (non-divergent) trait loci, while the pink line represents the mean FST in loci that underly trait divergence. Migration rates are *m* = 0 (A-B), *m* = 0.01 (C-D), *m* = 0.05 (E-F), *m* = 0.1 (G-H), *m* = 0.2 (I-J).

**Figure S4:**
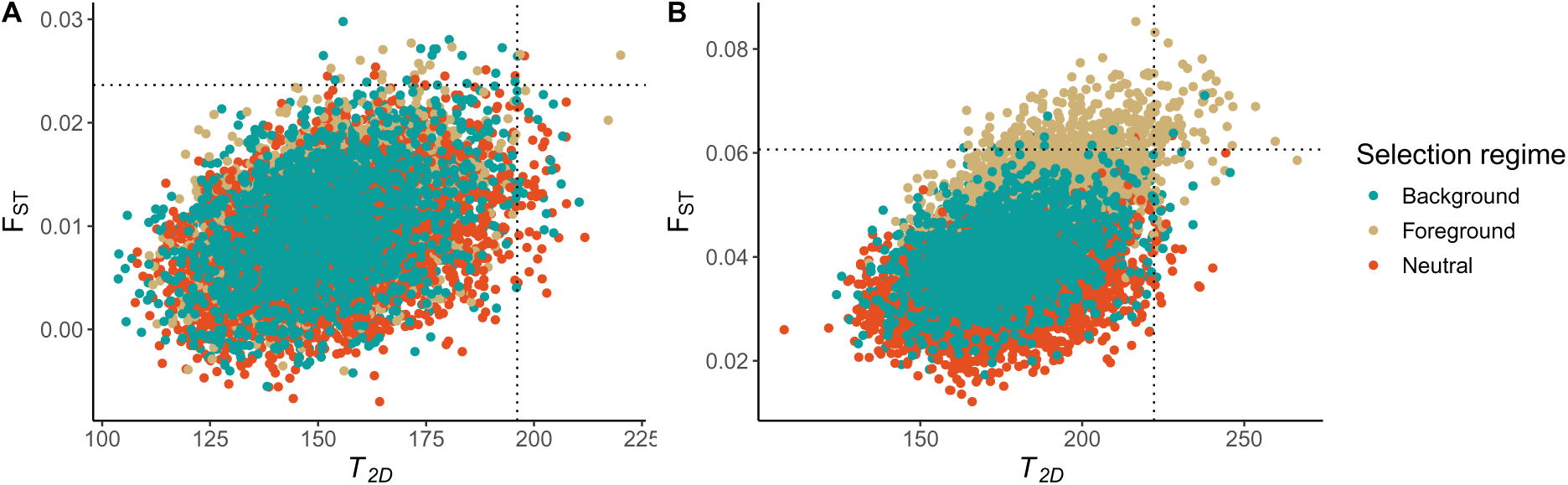
Comparison of *F_ST_* and *T*_2_*_D_* values at *t* = 300. (**A**) and *t* = 600 (**B**) for the CV model. The dashed lines represent the *α* = 5 *×* 10*^−^*^3^ threshold, as computed using the background loci.

**Figure S5:**
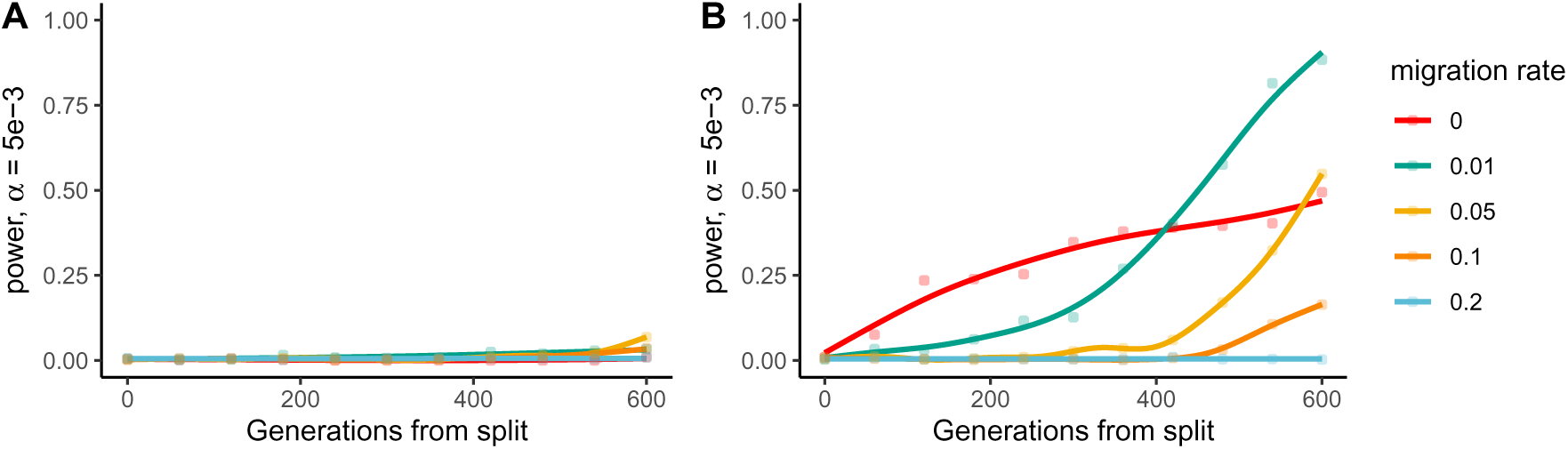
Power at *α* = 5 *×* 10*^−^*^3^ for the CV model for *T*_2_*_D_*. (**A**) and *F_ST_* (**B**), using selected loci to calculate the background distribution. Each point represents the fraction of loci undergoing divergent selection that had test statistics in the (1-*α*) right tail of the background test statistic distribution. Colors correspond to various migration rates as indiacated in the legend.

**Figure S6:**
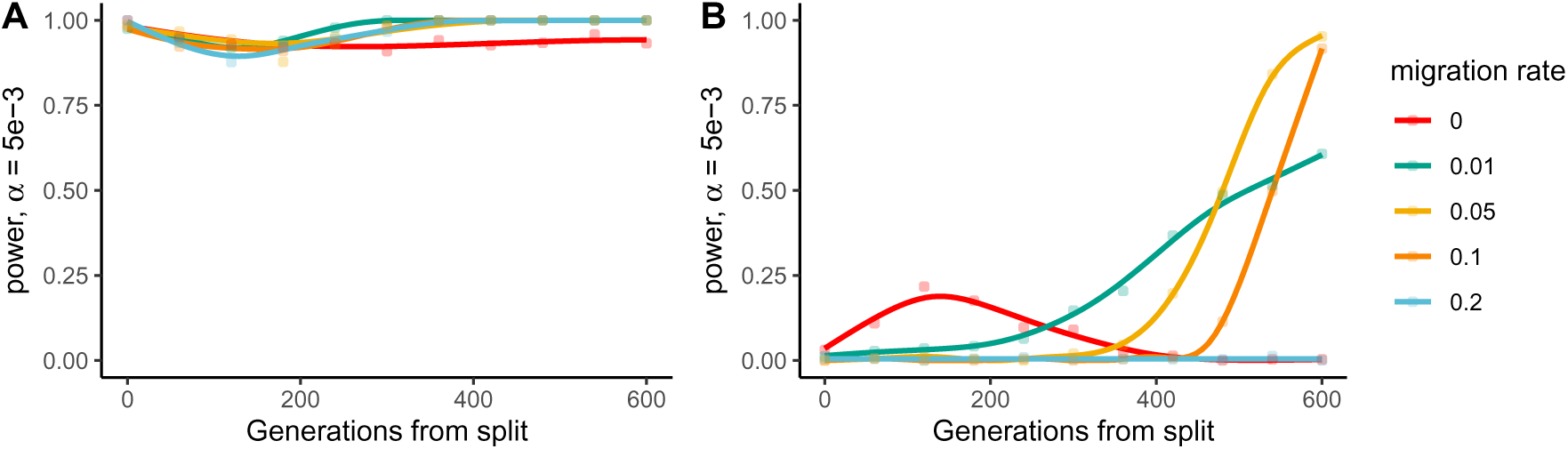
Power at *α* = 5 *×* 10*^−^*^3^ for the RV model for *T*_2_*_D_*. (**A**) and *F_ST_* (**B**), using neutral loci to calculate the background distribution. Each point represents the fraction of loci undergoing divergent selection that had test statistics in the (1-*α*) right tail of the background test statistic distribution. Colors correspond to various migration rates as indiacated in the legend.

**Figure S7:**
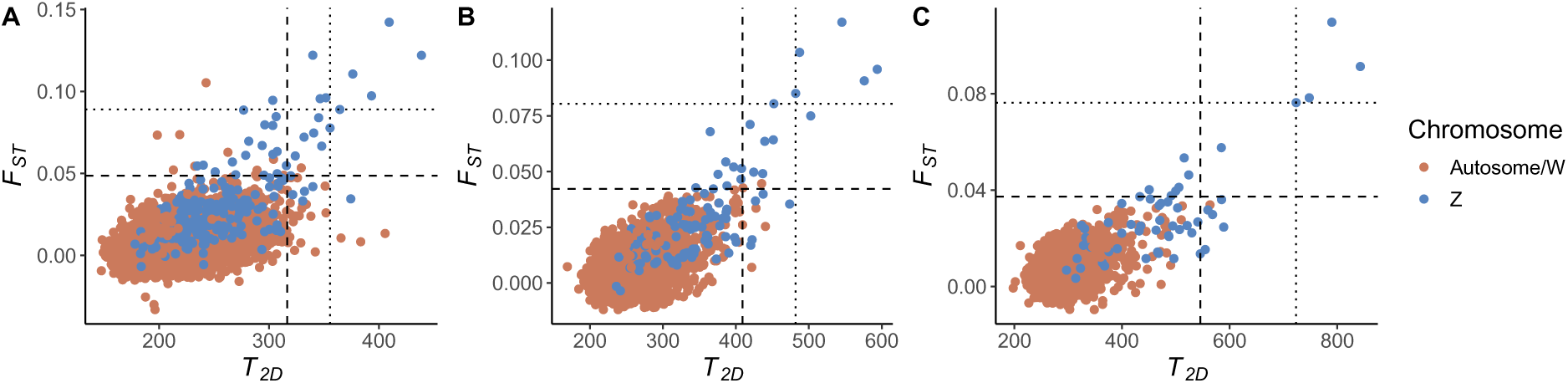
Comparison of *T*_2_*_D_* and *F_ST_* values in ECB genomic data. Each subpanel corresponds to a different window size (250 SNPs in **A**, 500 in **B**, and 1,000 in **C**). Chromosome Z, which was previously identified as harboring *F_ST_* outliers, is shown separately from the autosomes and the W chromosome. The dashed line represents the *α* = 5 *×* 10*^−^*^3^ threshold, while the dotted line represents the *α* = 1*e −* 3 threshold.

**Figure S8:**
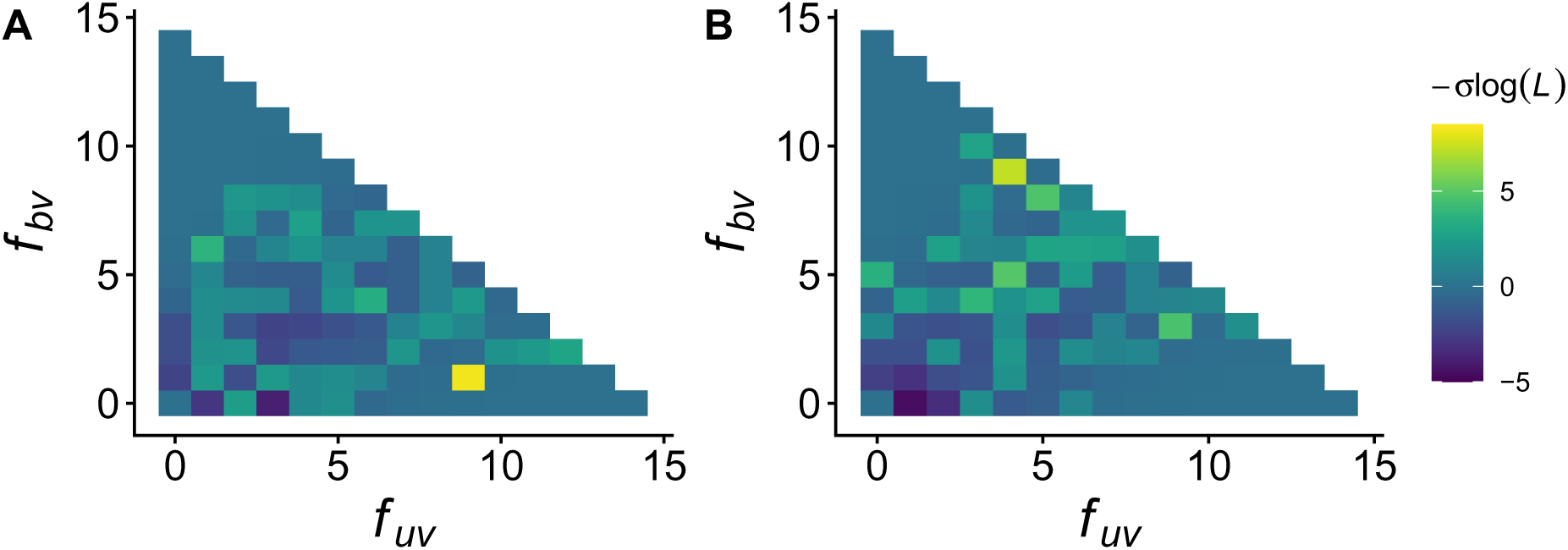
Likelihood calculations for two candidate selected regions containing *prl-1* and *rho1*. Since *T*_2_*_D_* is computed in frequency bins, we computed the relative contribution of each bin to the value of *T*_2_*_D_* in these loci. Colors represent the magnitude of the contribution to *T*_2_*_D_*, with positive values representing an excess of sites in the foreground and negative values representing an excess in the background.

**Figure S9:**
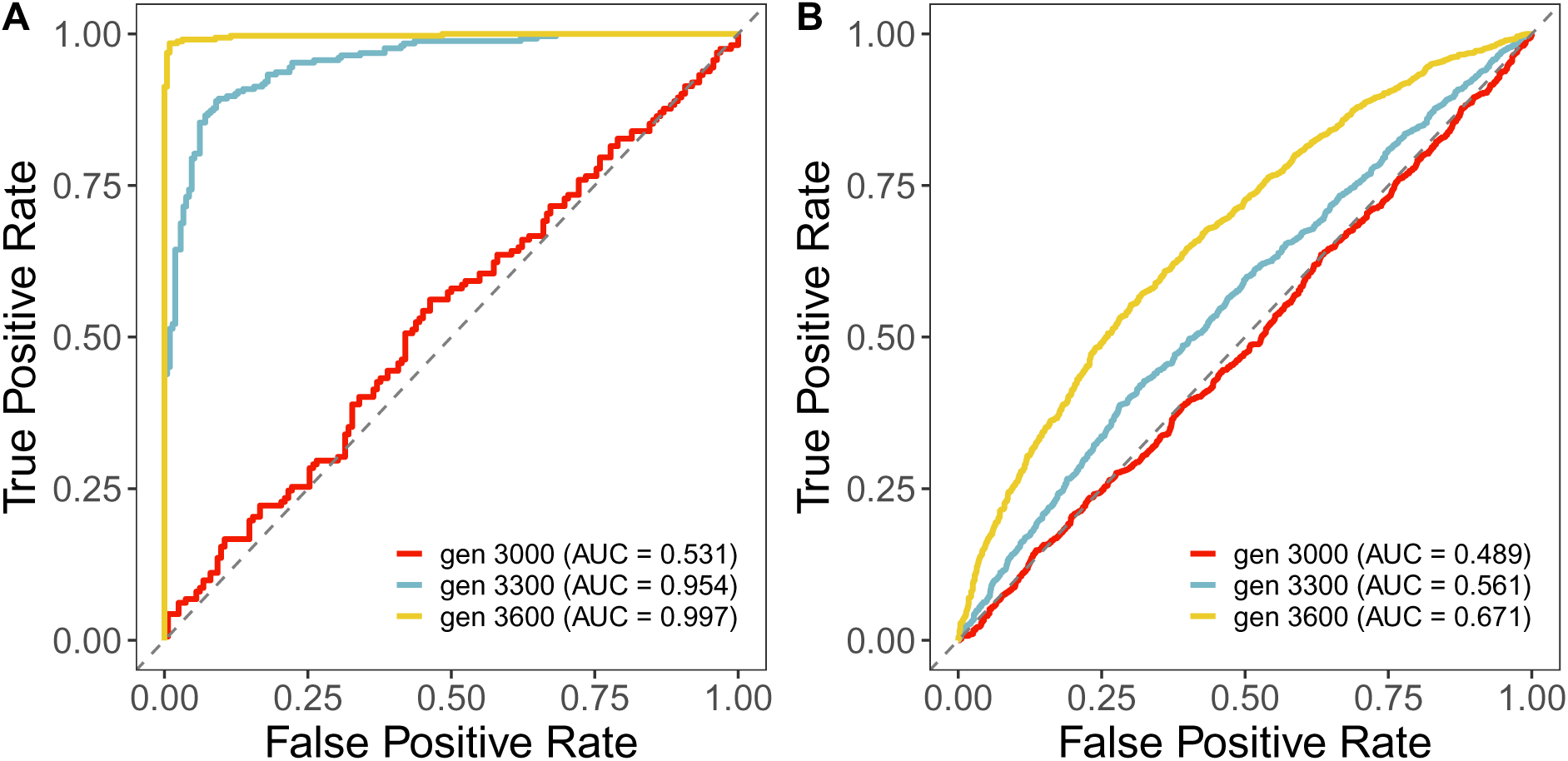
ROC curves for evaluation of the ability of *T*_2_*_D_* to distinguish windows under selection from neutral windows, using simulations with a migration rate of 0.01 from. **A:** the RV model and **B:** the CV model, for three time points across the simulation.

**Table S1:** Pearson correlations between *T*_2_*_D_* and *F_ST_* for windows in autosome/W and chromosome.

| Window size (number of SNPs) | Autosome/W |  | Z |  |
| --- | --- | --- | --- | --- |
| | $\rho$ | p-value | $\rho$ | p-value |
| 250 | 0.33 | $< 2.2 \times 10^{-16}$ | 0.74 | $< 2.2 \times 10^{-16}$ |
| 500 | 0.30 | $< 2.2 \times 10^{-16}$ | 0.80 | $< 2.2 \times 10^{-16}$ |
| 1,000 | 0.19 | $< 2.2 \times 10^{-16}$ | 0.80 | $1.563 \times 10^{-13}$ |

**Figure S10:**
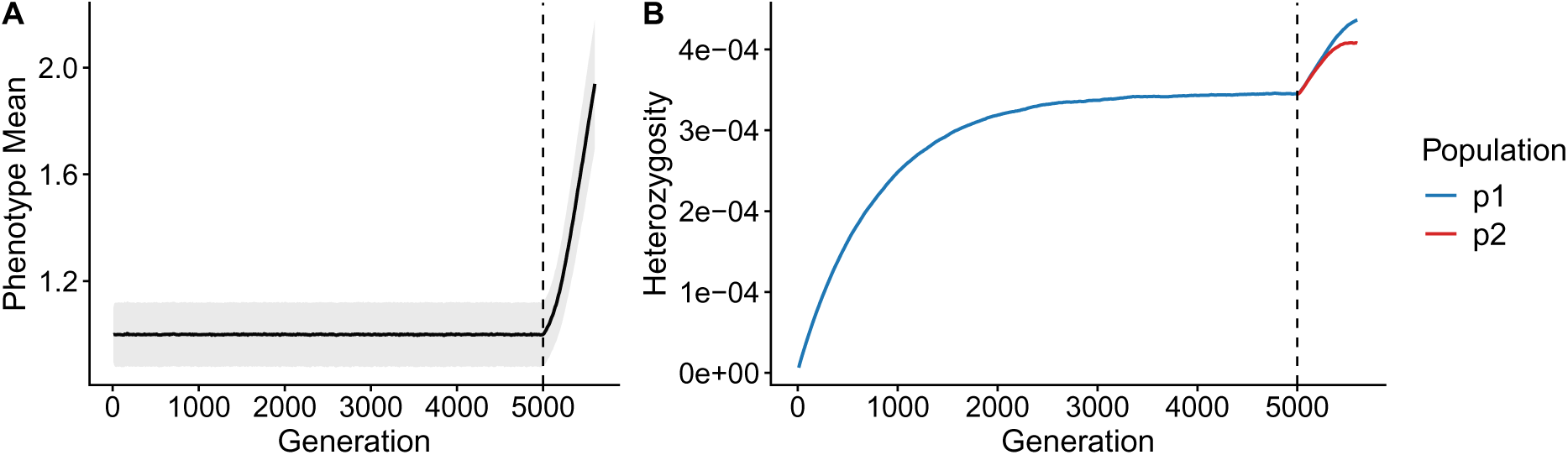
RV model simulations with 10*N* burn-in. **A**) Mean phenotype value over time with standard deviation represented by the grey envelope and **B**) mean heterozygosity estimated in both p1 and p2 subpopulations. Dashed lines represent the population split at *t* = 5000. Means were obtained by averaging over 100 simulation replicates.

**Figure S11:**
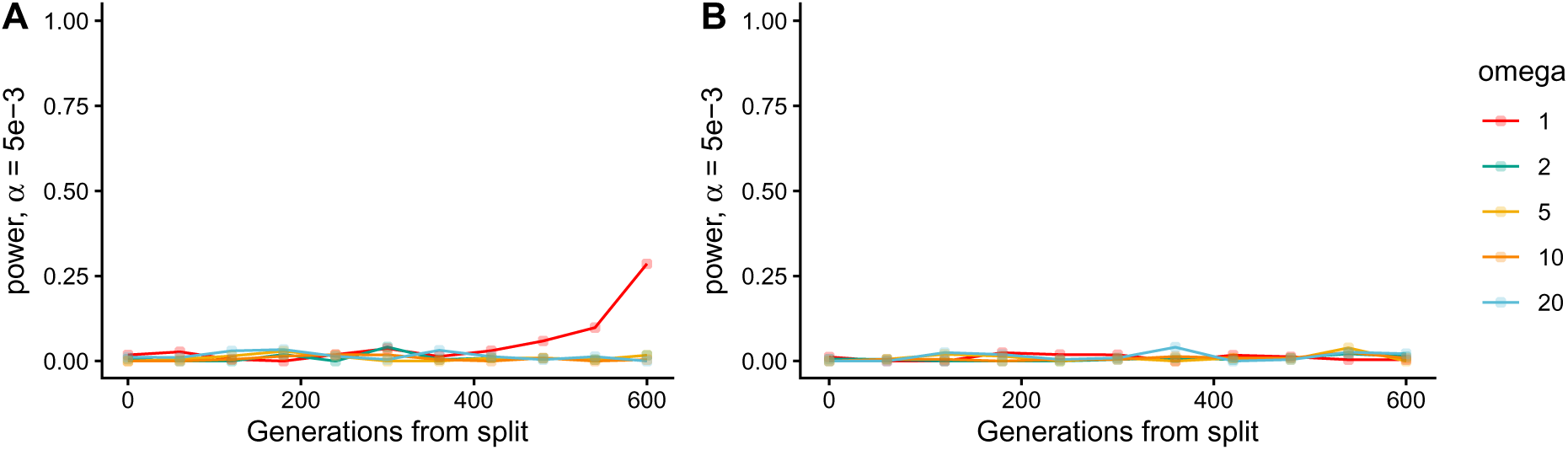
Power at *α* = 5 *×* 10*^−^*^3^ for the RV model for *T*_2_*_D_*. (**A**) and *F_ST_* (**B**) as a function of selection strength. All parameter values are the same as in the RV model presented in the main text. Each point represents the fraction of loci undergoing divergent selection that had test statistics in the (1-*α*) right tail of the background test statistic distribution. Colors correspond to various values of selection strength, and the solid lines represent a binomial regression on the data.

**Figure S12:**
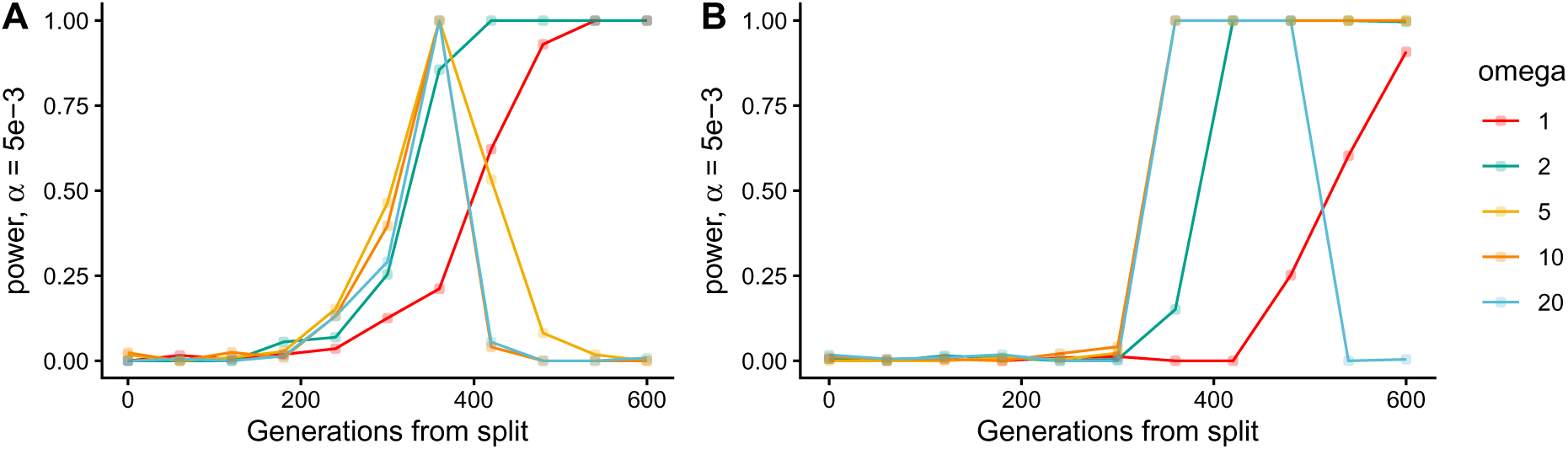
Power at *α* = 5 *×* 10*^−^*^3^ for the RV model for *T*_2_*_D_*. (**A**) and *F_ST_* (**B**) as a function of selection strength. All parameter values are the same as in the RV model presented in the main text, except that we increase the rate of environental change by a factor of *ω*^2^ relative to the value in Table 1. Each point represents the fraction of loci undergoing divergent selection that had test statistics in the (1-*α*) right tail of the background test statistic distribution. Colors correspond to various values of selection strength, and the solid lines represent a binomial regression on the data.

